# Predicting chromosomal compartments directly from the nucleotide sequence with DNA-DDA

**DOI:** 10.1101/2022.09.12.507578

**Authors:** Xenia Lainscsek, Leila Taher

## Abstract

3D genome architecture is characterized by multi-scale patterns and plays an essential role in gene regulation. Chromatin conformation capturing experiments have revealed many properties underlying 3D genome architecture such as the compartmentalization of chromatin based on transcriptional states. However, they are complex, costly, and time consuming, and therefore only a limited number of cell types have been examined using these techniques. Increasing effort is being directed towards deriving computational methods that can predict chromatin conformation and associated structures. Here we present DNA-DDA, a purely sequence-based method based on chaos theory to predict genome-wide A and B compartments. We show that DNA-DDA models derived from a 20 Mb sequence are sufficient to predict genome wide compartmentalization at the scale of 100 kb in four different cell types. Although this is a proof-of-concept study, our method shows promise in elucidating the mechanisms responsible for genome folding as well as modeling the impact of genetic variation on 3D genome architecture and the processes regulated thereby.

**Availability:** https://github.com/xX3N1A/DNA-DDA

**Supplementary information:** Supplementary data are available at … online.

## Introduction

Three-dimensional (3D) genome architecture allows linearly distal genomic loci to interact with one another, thereby impacting genome function. Chromosome conformation capturing techniques, in particular high-throughput chromosome conformation capture (Hi-C) [1, 2], have enabled us to systematically catalog genomic interactions and features of the 3D genome architecture in various cell types.

Hi-C data are typically summarized in a contact map, a matrix that estimates the probability of interaction between any two loci in the genome. Such maps are characterized by a plaid pattern reflecting enrichment or depletion of Hi-C interactions. This was observed already by early Hi-C studies, which proposed to segregate the loci into two sets of compartments, and arbitrarily termed them “A” and “B” [1]. Loci in A compartments preferentially interact with other loci in A compartments, while loci in B compartments tend to interact only with other loci in B compartments. Additionally, loci in A compartments are associated with transcriptionally active euchromatin, are gene-rich, and are centrally located in the nucleus [3]. In contrast, loci in B compartments are in transcriptionally inactive heterochromatin, and tend to be gene-poor and occupy the nuclear periphery [3]. A and B compartments have been shown to be associated with distinct histone acetylation and methylation patterns that reflect their transcriptional activity, and more refined subcompartmentalizations have been suggested on the basis of the observed chromatin states [2–5]. Compartmentalization has been found to be evolutionary conserved across species [6–8]. Nevertheless, it can differ substantially between cell types [3–5] and sequence variation between individuals has also been shown to underlie changes in 3D genome architecture, in many cases with pathological consequences [9–11].

Hi-C assays have revolutionized our understanding of 3D genome architecture, but they are expensive, time-consuming and expertise-demanding. Therefore, Hi-C data are only available for a limited number of human cells, and substantial effort has been put into deriving predictive computational models. Here we present DNA-DDA, a computational method that is based on the principles of chaos, ergodic and embedding theory to predict A/B compartments from the DNA sequence alone.

Chaos is widespread in biological systems [12–14]. It manifests itself as the seemingly random behavior of a deterministic process which is hypersensitive to fluctuations in initial conditions [15]. A deterministic dynamical system can be described by its current *state* (i.e., the system variables’ current values) and a system of differential equations (i.e., *rules*) which govern the evolution of the system (i.e., the sequence of states it passes through) [16, 17]. The set of all possible states, i.e., the solutions to the system of differential equations for every possible set of initial conditions, is known as the system’s *state space* [18, 19]. A *trajectory* in the state space is a sequence of states resulting from a particular set of initial conditions. For most chaotic systems, there exists an “attractor” [20, 21], i.e., a point or a set of points towards which trajectories from *almost any* set of initial conditions will approach, and that represents the long-term behavior –the dynamics– of the underlying system. Two initially infinitesimally close trajectories of such a system diverge exponentially and yet, bounded by their attractors, they will be similar in a topological sense; i.e., they can be deformed into each other continuously by stretching and folding [22]. Due to this counterintuitive property of “deterministic chaos”, the global structure of the state space can be investigated temporally, e.g., by studying the rate at which neighboring trajectories with similar initial conditions diverge as the system evolves [23], or spatially, e.g., by determining a trajectorie’s fractal dimension [24]. A bridge between these two perspectives is ergodic theory [25–27], which studies the statistical properties of a dynamical system. Trajectories of an ergodic system will eventually cover the entire state space so that under certain conditions, the time average of a function along a trajectory is related to the space average for almost all initial conditions [18].

The analysis of DNA sequences in the context of nonlinear dynamics (ergodic theory [28, 29] and chaos theory [14]) information theory [30, 31] and time series analysis (signal processing spectral analysis [32–34]) has a long standing history and their concepts and ideas are interconnected [35]. The aim of these methods is to study genomic control systems in relation with the laws of physics that govern their behavior. Inspired by many of the aforementioned concepts and ideas, we adapted a nonlinear time series classification framework, delay differential analysis (DDA) [36], for the prediction of A/B compartments from the DNA sequence.

The fundament of DDA is given by Takens embedding theorem, which states that under certain conditions, the measurement of a single variable of a high dimensional dynamical system providing a good or a global observability of the state space, can be sufficient to reconstruct the original state space [37, 38]. DDA relates delay and derivative embeddings of a variable in a (nonlinear) functional form, and uses the fitting coefficients as classifying features. A particular flavor of DDA, dynamical ergodicity-DDA (DE-DDA) [39], defines a measure for assessing dynamical similarity. We hypothesize that the DNA sequence and the interaction frequencies between genomic loci obtained from a Hi-C assay, are highly observable of 3D genome architecture, and that sequences in close proximity in 3D space will share certain dynamical properties. The Hi-C map can be understood as a 2D projection of the n-dimensional state space of the system, i.e., a recurrence plot [40]. A recurrence plot is a method from nonlinear data analysis obtained by recording the instances *t*, when a trajectory visits the immediate proximity of a state it has visited in the past. Analogously, a Hi-C map visualizes how often the genomic locus at position *t* is involved in interactions with a subsequent locus in the DNA sequence (i.e., how often it is revisited). The patterns which arise in the Hi-C maps are a hallmark for an underlying chaotic process.DDA has been applied most extensively in epilepsy research [41–43] and our study confirms its potential to be extended into the field of genomics. Therefore, inspired by the ergodic hypothesis, we examine the ensemble and time averages of dynamical information inherent in the numerical representation of the DNA sequences as described by DE-DDA, to infer their proximity in 3D space and, in turn, to predict A/B compartments.

## Methods

### Pre-processing of Hi-C data sequencing data

Raw FASTQ files from Hi-C experiments involving four cell lines were downloaded from the Gene Expression Omnibus database (Table 1; [2, 44, 47]) and mapped to the human reference genome (GRCh38/hg38) using `bowtie 2` (v.2.4.1; [49]) with options `—reorder and —very-sensitive-local`. The deepest sequenced data set which we considered was the “primary” GM12878 Hi-C data set comprising 3.6 billion sequence reads followed by the the K526 Hi-C data set with a library of 1.4 billion sequence reads. The older hESC and IMR90 data sets comprised 0.3 and 0.4 billion reads respectively.

**Table 1.** DNA-DDA structure selection and testing data sets.

| Cell type | Hi-C data | ChIP-seq | RS | DS |
| --- | --- | --- | --- | --- |
| GM12878 | GSE63525 [2], [44]<br>(primary) | GSE29611; [45]<br>GSM733772 | GATC | GATC |
| K562 | GSE63525 [2], [44] | GSE31755; [46]<br>GSM788085 | GATC | GATC |
| hESC | GSE35156 [47] | GSE29611; [45]<br>GSM733687 | AAGCTT | GATC |
| IMR90 | GSE35156 [47] | GSE103589; [48]<br>GSM2775001 | AAGCTT | GATC |

ChIP-seq: of histone mark H3K4me1 (GM12878, K562, IMR90) or H4K20me1 (hESC); RS: restriction sequence; DS: dangling sequence.

### Hi-C contact maps

The contact map of each autosome was generated from the mapped reads using the *HiCExplorer* pipeline (v.3.7.2; [50–52]). `hicBuildMatrix` was called with parameters “`—binSize 5000 —minMappingQuality 10 —restrictionSequence RS —danglingSequence DS`”, where DS and RS are the restriction and dangling sequences listed in Table 1. The resulting Hi-C matrix was balanced with the algorithm introduced by Knight and Ruiz [53] using `hicCorrectMatrix correct`; the “`—filterThreshold`” parameter was chosen based on the histogram produced by `hicCorrectMatrix diagnostic_plot` (Supplementary Table S1). From this matrix, a contact map at the resolution of 100 kb was derived using `hicMergeMatrixBins` with parameter “`--numBins 20`”.

For comparability, the contact map obtained in this manner was scaled to the 0 to 1 range with `hicNormalize --normalize normrange`. An entry in the resulting matrix **H** = (*h_i,j_*) represents the contact frequency between genomic bins *i* and *j*. Bins enclosing contromere locations (obtained from the UCSC table browser [46]) as well as low coverage bins (below 10% of overall interaction probability) were excluded from analyses (Supplementary Tables S5 -S8).

### DNA-DDA

#### Delay differential analysis in general

Let *x*(*t*) be a dynamical sequence of length *L* where *t* represents increments in time or space. A nonlinear DDA model has the following general functional form ([39] and citations therein):

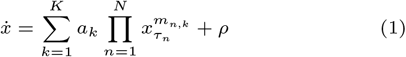

where 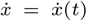 is the derivative of the original time (or space) series *x* = *x*(*t*) and *x*(*t* — *τ*) is the value of the series shifted by *τ* steps *x_τ_n__* = *x*(*t* — *τ_n_*). The model parameters are: *K*, the number of monomials; *N*, the number of delay embeddings *x_τ_n__* contained in each monomial; *m_n,k_* ∈ **N**_0_, the order of nonlinearity of the nth delay embedding in the *k*th monomial. Finally *a*_1_, *a*_2_, *a*_3_ are the fitting coefficients, and *ρ* is the least square error of the model:

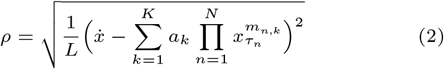

A full list of all possible three term DDA models up to cubic nonlinearity and two delay pairs (*K* ∈ {1,2, 3}, *m* ∈ {1,2, 3}, *n* ∈ {1,2}) can be found in Supplementary Table S4.

For a dynamical sequence *x*(*t*), Equation 1 can be written as the over-determined system of equations

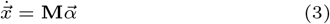

where each element in 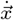 is the center derivative at each time/space point 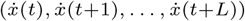, **M** a matrix where each column represents a monomial (delay embedding 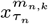) at a certain time/space point (row), and 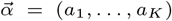 are the fitting coefficients of the model which are estimated for the input data using singular value decomposition (SVD), and together with the least square error (Equation 2) make up the classifying feature set 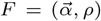. Furthermore, Equation 1 can be solved for *I* dynamical input sequences 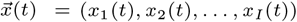 either individually (single-trial (ST) DDA) or simultaneously (cross trial (CT) DDA) (Figure 1) by extending the over-determined system of equations of Equation 3 to

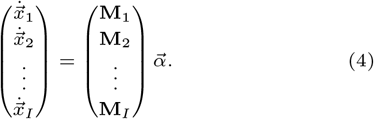

**Fig. 1.**
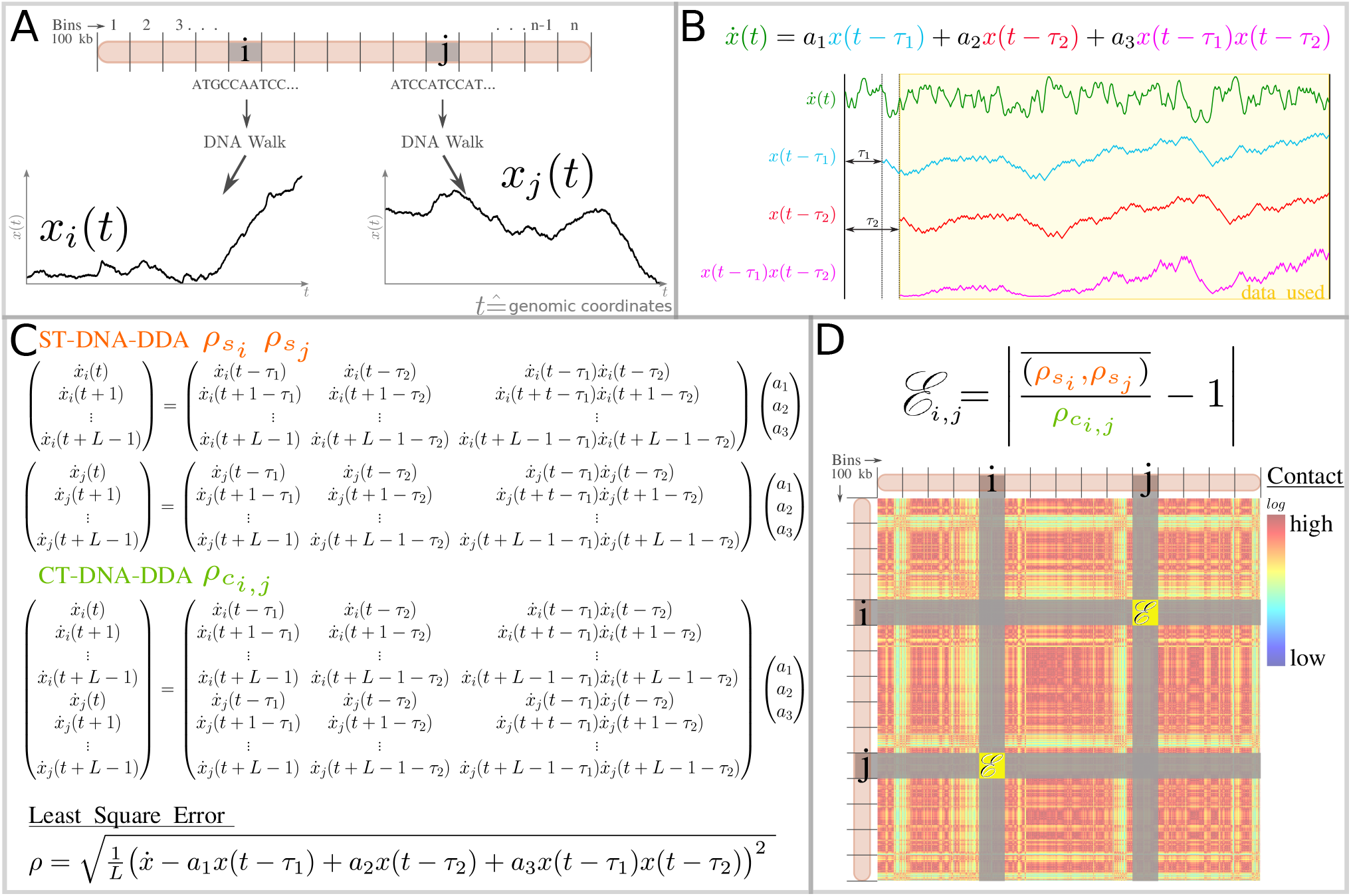
Steps taken to derive the entries of a DNA-DDA contact matrix D. A) The DNA sequences of two exemplary bins *i* and *j* located on one chromosome are represented as *DNA walks x_i_*(*t*) and *x_j_*(*t*). Each walk starts at *x*(*t* = 1) = 0 and takes a step up (+1) if the nucleotide at position *t* + 1 is C or G and down (−1) if the nucleotide at position *t* + 1 is A or T. B) The DNA-DDA model and visualization of one DNA walk *x*(*t*) and its respective derivative 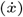 and delay embeddings (*x*(*t* — *τ*_1_) and (*t* — *τ*_2_). The yellow window indicates the data points used in estimation of the DDA features (*a*_1_, *a*_2_, *a*_3_, *ρ*). The yellow window indicates the data points used in the estimation of the DDA features (*a*_1_, *a*_2_, *a*_3_). C) Overdetermined system of equations for ST and CT DDA. The coefficients (*a*_1_, *a*_2_, *a*_3_) are determined by singular value decomposition (SVD) separately for bin *i* and *j* in ST DDA and in a single step in CT-DDA. Least square errors are computed for ST DDA (*ρ_s_i__*, *ρ_s_j__*) and CT DDA (*ρ_c_i,j__*). D) The ST DDA least square errors (*ρ_s_i__*, *ρ_s_j__*) and CT DDA least square error *(p_Ci_**,j***) are combined to dynamical ergodicity 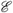 [39]. Note this figure is a detailed description of how the DNA-DDA matrix is determined in the workflow of DNA-DDA (see Supplementary Figure S3).

ST and CT DDA features have recently been combined by Lainscsek et. al [39] in a way which allows testing for dynamical similarity between two dynamical input sequences *x_i_*(*t*) and *x_j_*(*t*). Here the mean of the ST error is representative of the temporal average, and the CT error is representative of the ensemble average. In accordance with the ergodic hypothesis, if *x_i_*(*t*) and *x_j_*(*t*) are dynamically similar, the mean of the ST errors 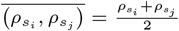 and the CT error (*ρ_c_i,j__*) will also be similar and therefore, their quotient close to one. Dynamical ergodicity (DE) DDA 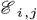 is defined as

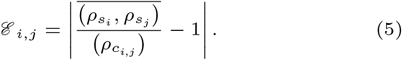

The smaller 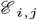, the more dynamical similar the two signals under investigation, *i* and *j*, are.

#### DDA for genomic sequence data

##### Numeric representation of DNA sequences

For DNA-DDA, *t* corresponds to the genomic position of a nucleotide and *L* is the resolution or number of nucleotides in each bin. We modeled the DNA sequence as a 1D random walk (DNA walk). Specifically, for every genomic bin, the walker starts at *x*(*t* = 1) = 0 and progresses along the DNA sequence taking a step upwards (*x*(*t* + 1) = *x*(*t*) + 1) if the nucleotide at position *t* + 1 is C or G and down (*x*(*t* + 1) = *x*(*t*) — 1) if the nucleotide at position *t* + 1 is A or T [54]. The DNA walk of five exemplary nonempty bins covering the genomic region chr1:800001-1300001 is depicted in Supplementary Figure S1. A time delay in a DNA-DDA model is a shift in genomic coordinates. A delay embedding of the DNA sequence relates the value of the DNA walk at the genomic coordinate *t* to its “previous” value at *x*(*t* — *τ*). We are trying to gain access to (1) the interaction frequency by considering only (2) the DNA nucleotide sequence. We achieve this by exploiting the concept of embedding theory, which states that certain variables of nonlinear systems are coupled and entail information about one another.

##### DE-DDA classifying feature set computation

The sequence of the GRCh38/hg38 assembly of the human genome was partitioned into 100 kb non-overlapping bins, and the sequence of each bin represented as a 1D DNA walk. For a pair of bins *i,j*, we computed the ST- and CT-errors *ρ_s_i__*, *ρ_s_j__* and *ρ_c_i,j__* using a C implementation of DDA provided by the author [39]. This executable takes as input a time series and parameters listed in Supplementary Table S2 and outputs ST- and CT-classifying feature set (αi,α2,α3,p). We combined the errors *ρ_s_i__*, *ρ_s_j__* and *ρ_c_i,j__* into 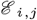 (Equation 5), which we propose as an estimation of the contact probability between the two pairs of genomic bins. We repeated this process for all bin pairs to obtain the DNA-DDA contact map **D** = (*d_i,j_*) of a given chromosome.

##### Matrix Post-processing

We hypothesize that bins with higher interaction probability will be similar in certain dynamical properties. Thus, we inverted **D** so that highest values are mapped to the lowest and vice versa. The logarithm of each non-zero value in **D** was taken. The matrices were saved in the file format of `HOMER` [55] and then converted to h5 with *HiCExplorer*’s `hicConvertFormat` function. The resulting matrices were normalized to the 0 to 1 range with `hicNormalize —normalize normxange`. Bins that were excluded in the Hi-C contact maps **H** were also excluded in the DNA-DDA contact maps **D**.

### Compartment calling

To call compartments, we first derived the Pearson correlation matrix **C_H_** = (*c_H__i,j_*) from the normalized Hi-C matrix **H** as described by Lieberman, and then applied principal component analysis (PCA) to **C_H_** using MATLAB®’s `pca` function. For each PC, values larger than three scaled median absolute deviations (MAD) from the median were considered extreme outliers and replaced with nearest value that was not an outlier using MATLAB®’s `filloutliers(PC,’nearest’)` function. The PCs were then normalized to zero mean and unit variance. Lastly, values greater 0 were assigned to the A compartment and scaled to [0, 0.5]; values below 0 were assigned to the B compartment and scaled to [0.5, 1]. We used ChIP-seq data for H3K4me1 or H4K20me1, two epigenetic modifications associated with open chromatin [56], to determine which principal component (PC) defines compartments (Table 1). More specifically, among the first four PCs, we selected the one with the largest absolute Pearson correlation coefficient with the ChIP-seq profile to be representative of A/B compartments (Supplementary Table S9). ChIP-seq profiles were generated from the corresponding bed files; an empty vector of the same length as the PC was generated and +1 was added to for each called peak falling into the genomic region of a particular bin. If the Pearson correlation coefficient was negative, the PC was multiplied by −1. In two exceptional instances (chr1 for Hi-C contact maps of GM12878 and IMR90), a different PC with approximately the same Pearson correlation coefficient was deemed more likely to be associated with the compartments upon inspection, and used instead. From here on, we refer to the PC obtained from **C_H_** as PC_Hi-C_.

Computationally derived DNA-DDA matrices **D** were processed in the same manner with one exception, PC_DNA-DDA_ were smoothed with a sliding window of 5 using MATLAB®’s `movmean()` function before computing the Pearson correlation coefficients with the ChIP-seq profiles. We further refer to the resulting DNA-DDA Pearson correlation matrices as **C_D_** = (*c_D__i,j_*) and to the corresponding PCs defining A/B compartments as PC_DNA-DDA_.

#### Saddle plot analysis

We performed saddle plot analysis on the contact maps predicted by DNA-DDA and Hi-C respectively in the K562 and IMR90 cell lines. For each chromosome, values of P_Hi-C_ and PC_DNA-DDA_ were split into 30 equisized bins (PC_Hi-C-S_ and PC_DNA-DDA-S_). The expected interaction value at distance *d* of the balanced [53] Hi-C contact map **H** was computed as the sum of its *d*th diagonal *diag_d_*(**H**), divided by the number of elements in diagonal *d*, *n*(*diag_d_*(**H**)). Then we sorted the values of the expected/observed Hi-C matrix **H**_*oe*_ and the DNA-DDA matrices **D** according to PC_Hi-C-S_ and PC_DNA-DDA-S_ respectively. Finally, we quantified compartment strength from the interaction values that were allocated to the highest 25% of the PC values (implying *AA* and *BB* interactions) and lowest 25% of PC values (implying *AB* or *BA* interactions) as 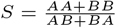. Note that all PC values were scaled to the interval [—1, 1], as often done (e.g., in [4]), and not as explained in the *Compartment calling* section, as this would have resulted in a large number of bins with no allocated PC values.

#### Correlation with CG content

We compared DNA-DDA’s performance to predict compartments with a baseline of AT/CG percentage over 100 kb windows in all four cell types.

### Structure selection and testing

A key difference to traditional machine learning-based approaches is that DDA models are not updated iteratively (i.e., they do not learn). Instead, an exhaustive sweep is performed over a list of putative models to search for those best suited to discriminate the dynamics of interest; this step is called *structure selection*.

The functional form of a DDA model is dictated by the overall system and obtained data type. The data type most extensively studied using DDA is EEG, for which a particular functional form has been established [41–43, 57, 58]. Typically, most terms in Equation 1 are set to zero to reduce the chances of overfitting (e.g., *K* ∈ {1,2, 3}, *m* ∈ {1,2, 3} *n* ∈ {1,2}). This work is a proof of concept to explore the feasibility of applying DDA to genomics data. Thus, we decided to use a simple, symmetric model with only quadratic degree of nonlinearity (DDA model number 2 in Supplementary Table S4):

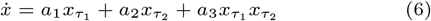

Symmetric models require half the computational effort of non-symmetric ones to compute the classifying feature set {*a*_1_, *a*_2_, *a*_3_, *ρ*} for a range of delays between *r*_1_ and *r*_2_; *τ*_1_, *τ*_2_ ∈ (*r*_1_, *r*_2_).

Next, we searched for the delay pair *τ*_1_, *τ*_2_ that best captured A/B compartments in each of the four cell-types. Specifically, this was done using a 20 Mb region on chr22 (chr22:16200000-36200001, Supplementary Figure S2). The chromosome was chosen arbitrarily and the region thereon corresponded to the one with the lowest compartment agreement between the four cell types considered in this study. The performance of each delay pair was measured as the Pearson correlation coefficient between PC_Hi-C_ and PC_DNA-DDA_ in this region.

The four obtained DNA-DDA models were tested on s all 100 kb genomic loci of all human autosomes with various performance measures (Pearson correlation coefficient *r_PC_* between PC_Hi-C_ and *PC_DNA-DDA_*, area under the roc curve `AUC`, accuracy `ACC` and F1-score `F1` for classifying A/B compartments). We would like to emphasize that once we determined the DNA-DDA model for each cell type (Supplementary Table S3), all subsequent analyses and results were performed on never before seen data (with the small exception of chr22:16200000-36200001).

### Comparison to other methods

We compared DNA-DDA to three other methods that can predict A/B compartments from sequence-based features or the sequence directly (Table 2). The “Sequence-based Annotator of chromosomal compartments by Stacked Artificial Neural Networks” (SACSANN [8]) predicts A/B compartments based on features derived from GC content, transposable elements (TE), and putative transcription factor binding sites (TFBS). It identifies the 100 most important species/cell-type specific features with a random forest predictor and then trains two stacked artificial neural networks (ANN) to classify 100 kb-long genomic bins into either the A or B compartment.

**Table 2.**
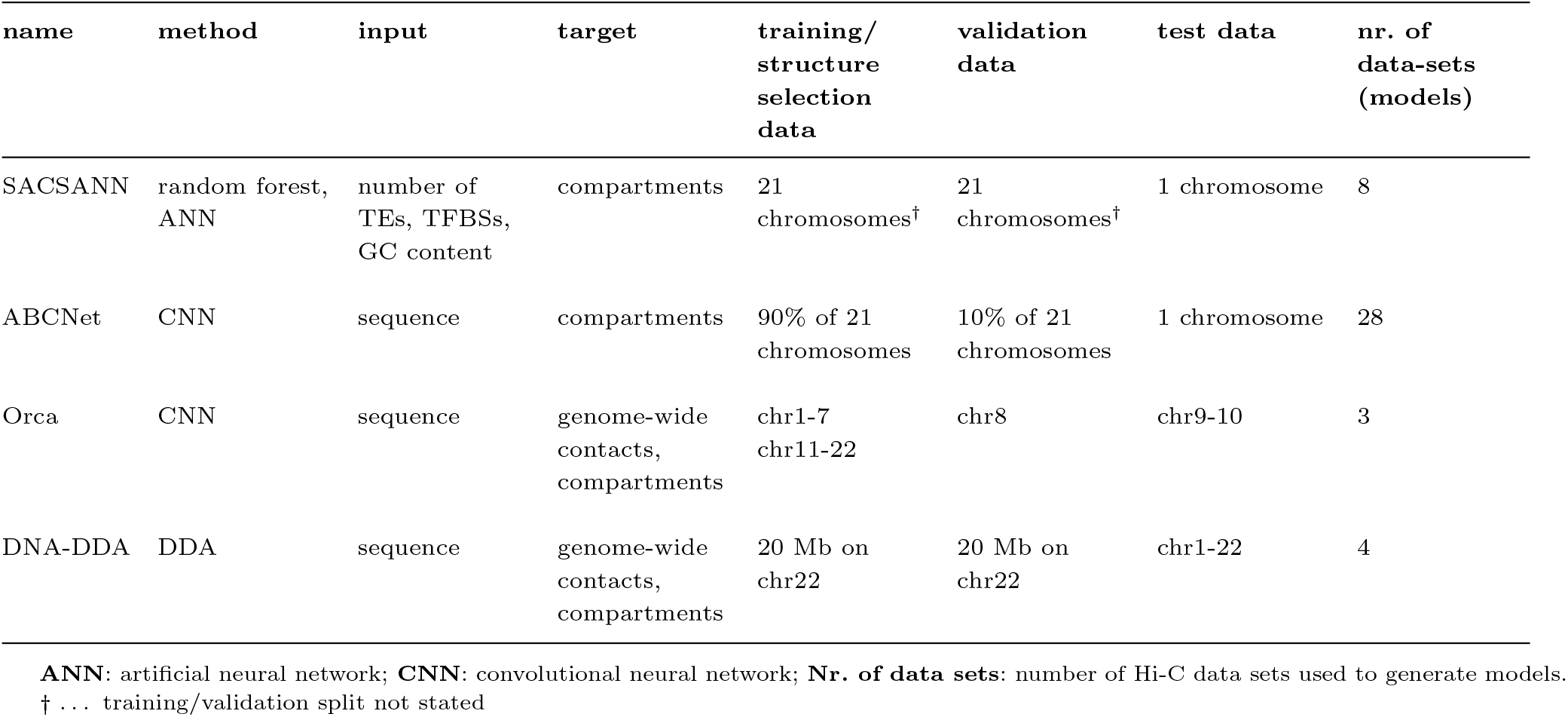
Comparison of DNA-DDA models to other methods

| name | method | input | target | training/<br>structure<br>selection<br>data | validation<br>data | test data | nr. of<br>data-sets<br>(models) |
| --- | --- | --- | --- | --- | --- | --- | --- |
| SACSANN | random forest,<br>ANN | number of<br>TEs, TFBSs,<br>GC content | compartments | 21<br>chromosomes<sup>†</sup> | 21<br>chromosomes<sup>†</sup> | 1 chromosome | 8 |
| ABCNet | CNN | sequence | compartments | 90% of 21<br>chromosomes | 10% of 21<br>chromosomes | 1 chromosome | 28 |
| Orca | CNN | sequence | genome-wide<br>contacts,<br>compartments | chr1-7<br>chr11-22 | chr8 | chr9-10 | 3 |
| DNA-DDA | DDA | sequence | genome-wide<br>contacts,<br>compartments | 20 Mb on<br>chr22 | 20 Mb on<br>chr22 | chr1-22 | 4 |

**ANN:** artificial neural network; **CNN:** convolutional neural network; **Nr. of data sets:** number of Hi-C data sets used to generate models.
<sup>†</sup> ... training/validation split not stated

The “A/B Compartment Network” (ABCNet [59]) is a deep convolutional artificial neural network (CNN) that takes a one-hot encoding of the sequence within 100 kp-long bins and extracts features by passing them through two-layer convolutional kernels, an average pooling layer, and a fully connected dense layer to output a single value representing the predicted PC value of each bin, classifying it either as an A or B compartment.

Finally, “Orca” [60] is a multiscale prediction model composed of two CNNs, a hierarchial multi-resolution sequence encoder and a cascading series of sequence decoders. The encoder takes up to 256 Mb one-hot-encoded sequence as input, and generates a series of decreasing resolution sequence representations, centered around the input, with a convolutional architecture. The decoders then each predict interactions of up to 256 Mb at 1024 kb resolution at the top level and interactions within 1 Mb at 4 kb resolution at the bottom level.

Both SACSANN and ABCNet model architecture were applied to numerous data sets (Table 2) in terms of feature selection, training/testing procedures. Resulting models were evaluated using a chromosome-wise leave-one-out cross validation. We compared the performance of DNA-DDA on the hESC data set (GSE35156) to that reported by SACSANN’s and ABCNet’s authors on the same data set. In addition, we compared the *overall* performance of the methods, i.e., the average *AUC* or *ACC* across various data sets and models. In the case of SACSANN, we considered all examined data sets (“Summary (ROC AUC score) SACSANN” in supplemental_file_S3.xls at https://github.com/BlanchetteLab/SACSANN/tree/master/supplemental_files/,last accessed in July 2022). For ABCNet we restricted the comparison to the *ACC* reported for human data sets (i.e., average *μ* across all “secondary” data sets in Table III in Kirchof [59].

Orca was trained on two of the highest resolution micro-C (an improvement of Hi-C) data sets available for the H1 human embryonic stem cell line (H1-ESC) and Human foreskin fibroblast cell line (HFF) [61]. We compared to Orca models which predict interactions at the closest resolution that DNA-DDA currently operates on (128 kb and 100 kb respectively). To increase comparability, we considered an Orca model trained on the same cell line (H1-ESC 4DNES21D8SP8); it takes 32 Mb as input and predicts 128 kb interactions within this region. Therefore, we split one of the Orca hold-out chromosomes, chr9, into three 32 Mb regions (Supplementary Tab. S11), derived a Pearson correlation matrix from the predicted Orca contact maps, performed a PCA in MATLAB® and computed the Pearson correlation coefficient r_PC_ of the resulting PCs to those identified by the HiC-data for hESC GSE35156 at 128 kb.

## Results

DNA-DDA is a statistical approach derived from DDA, a method based on nonlinear dynamics and traditionally used for time series data, which predicts A/B compartments from the reference sequence (Figure 1). DDA models relate the numerical derivatives of the input data to their time-delayed versions in a nonlinear functional form, of which the fitting coefficients and model error {*a*_1_, *a*_2_, *a*_3_, *ρ*} are used to access information about the underlying dynamical system. With simplicity and efficiency in mind, we chose the functional form given in Equation 6. In principle the analysis can be summarized in three steps: (1) segment the reference sequence into 100 kb long bins and represent the sequence in each bin as a 1D DNA walk, (2) determine the best suited model parameters (delay pair *τ*_1_, *τ*_2_) in each cell type by supervised structure selection with Hi-C derived compartment labels, and (3) apply the model to the rest of the genome to predict A/B compartments.

### Structure selection

To find the delay pairs *τ*_1_, *τ*_2_ in Equation 6 that best capture A/B compartments in each cell-type, we tested model performances for all possible combinations of values for *τ*_1_, *τ*_2_ on a 20 Mb-long region of chr22 (Methods). Briefly, we partitioned this region into 200 100 kb-long bins, estimated the interaction probability between each pair of bins *i* and *j* as 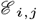 (Equation 5), built a DNA-DDA contact map **D**, apply PCA to the respective DNA-DDA Pearson correlation matrix **C_D_** to call A/B compartments. For each cell type, we then chose the delay pair that resulted in the highest absolute Pearson correlation coefficient r_PC_ between PC_DNA-DDA_ and PC_Hi-C_ (Supplementary Figure S4). This resulted in four different delay pairs, one for each cell type (Supplementary Table S3). Pearson correlation coefficients between PC_DNA-DDA_ and PC_Hi-C_ ranged between 0.21 (hESC) and 0.49 (GM12878) and AUCs ranged between 0.61 (hESC) and 0.77 (GM12878).

### Testing

The DNA-DDA Pearson correlation matrices **C_D_** exhibited strikingly similar global patterns to the experimentally obtained Hi-C Pearson correlation matrices **C_H_** (Figure 2 and Supplementary Figures S9 to S12). In general, supervised methods are limited by the uncertainty of the labels derived from the experimental data. It is known that identifying A and B compartments by PCA of the Pearson correlation matrix derived from the contact map is suboptimal and misses many features, especially at higher resolutions. This is particularly apparent in the IMR90 data set (Supplementary Figure S7) or particular chromosomes (e.g., chr22). Nevertheless, DNA-DDA achieves exceptional performances on never before seen genome, (with 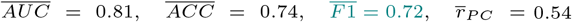 over all chromosomes and cell lines; Table 3), especially considering the mere 20 Mb region that was used to determine the model/delay pair combination. The model/delay pair combination achieves the highest performance in the K562 cell line (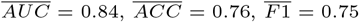, and 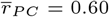), and the lowest performance in the IMR90 cell line (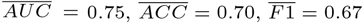, and 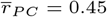).

**Fig. 2.**
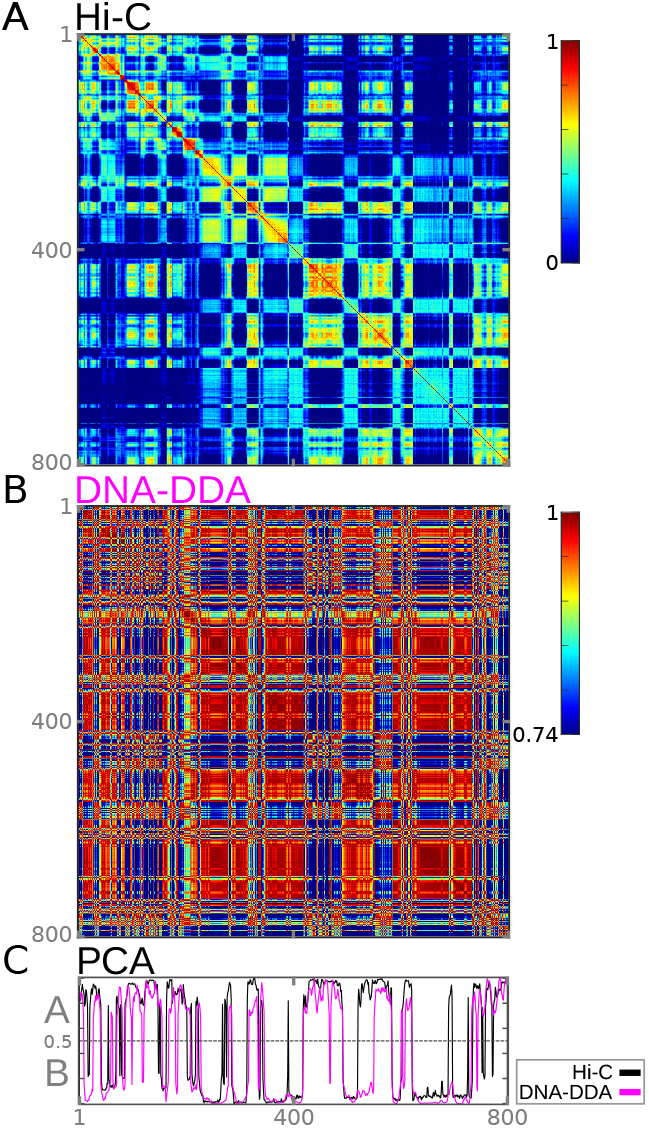
DNA-DDA predicts A/B compartments from the DNA sequence alone. A) Hi-C (C_H_) and B) DNA-DDA (C_D_) Pearson correlation matrices of the K562 cell line for an example hold out chr18 exhibit strikingly similar patterns. The color scale of the DNA-DDA Pearson matrix goes from its mean minus variance (C_D__*μ*_ — C_D__*σ*^2^_) to its maximum. C) Resulting PC_Hi-C_ (black) and PC_DNA-DDA_ (magenta) used to define A/B compartments are in very strong correlation to one another (*r_PC_* = 0.73).

**Table 3.**
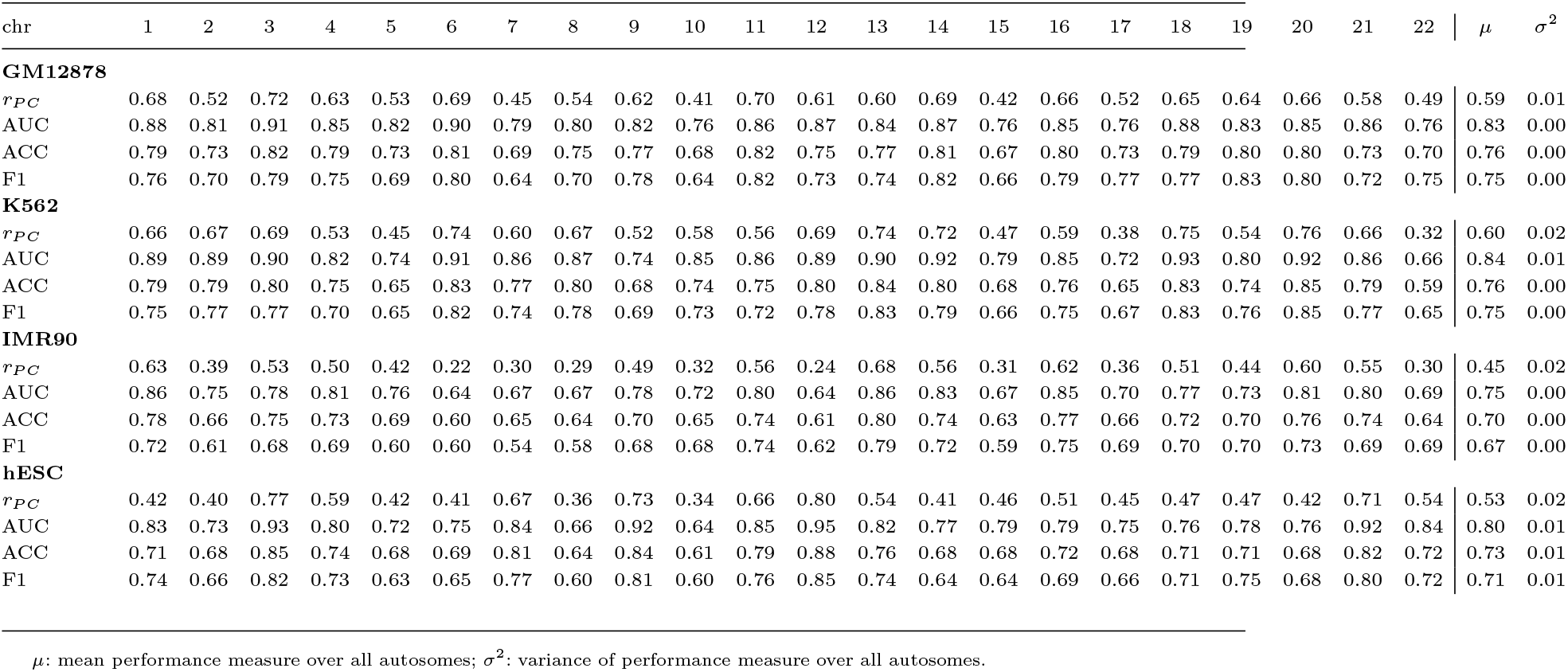
Performances of DNA-DDA

#### Saddle plot analysis

Saddle plot analysis revealed that DNA-DDA derived matrices had overall stronger saddle strengths strengths *S* than those derived from the expected/observed Hi-C matrix **H**_o*e*_ (Supplementary Figure S15). The mean saddle strength across all chromosomes for each cell line was 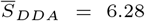 and 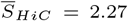 for K652, 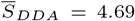 and 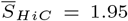 for IMR90 and 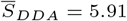 and 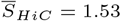 for hESC. However, this analysis should be interpreted with caution as the Hi-C and DNA-DDA contact maps are of very different origins.

#### Correlation with CG content

CG content is known to correlate with the compartment signal. Nonetheless, DNA-DDA predicted compartments better for 100 kb windows than the corresponding percentage of AT/CG content in each window. The mean Pearson correlation coefficients across all chromosomes was between 0.45 and 0.60 as opposed to the means between 0.45 and 0.56 observed for the GC-content-based prediction. Specifically, when examining individual chromosomes, DNA-DDA performed significantly better for GM12878, K562 and hESC (*p* < 0.05; Wilcoxon signed-rank test) and exhibited no difference for IMR90, the cell line for which DNA-DDA performed the worst overall. These results suggest that DNA-DDA and GC content are complementary methods.

### Comparison to other methods

We compared DNA-DDA to SACSANN [8], ABCNet [59] and Orca [60]. To our knowledge, these are the only methods that are directly comparable to DNA-DDA, as they rely only on the DNA sequence or sequence-based features to predict A/B compartments. DNA-DDA competes well with other methods predictive of A/B compartments when assessed on the same Hi-C data set (hESC GSE35156) as well as overall (Table 4). Indeed, all four methods showed similar scores measured by *AUC*, *ACC* or *r_PC_*.

**Table 4.** Comparison of DNA-DDA performance to other methods

| | Performance measure | hESC (GSE35156) | $\mu_1$ |
| --- | --- | --- | --- |
| SACSANN | AUC | 0.81 | 0.83 |
| DNA-DDA | AUC | 0.82 | 0.81 |
| ABCNet | ACC | 0.75 | 0.80 |
| DNA-DDA | ACC | 0.75 | 0.76 |
| | Performance measure | hESC (GSE35156) regions 1-3 | $\mu_2$ |
| Orca | $r_{PC}$ | 0.86, 0.87, 0.90 | 0.89 |
| DNA-DDA | $r_{PC}$ | 0.76, 0.71, 0.74 | 0.74 |

**hESC (GSE35156) regions 1-3:** Pearson correlation $r_{PC}$ of predicted PCs of DNA-DDA and Orca contact maps at 100 kb and 128 kb respectively with three 32 Mb regions on chr9 in hESC.
$\mu_2$ : mean prediction performance across the three 32 Mb regions.

Note the vast difference in the size of the “*training*” (pendant to structure selection in DNA-DDA) and test data sets between the methods in Table 2. Furthermore it should be noted that although the Orca model we compared to was trained on the same cell line, a far more deeply sequenced HiC library was used in the training/validation procedure (0.3 billion vs 5.6 billion (4DNES21D8SP8) total reads).

## Discussion and Conclusion

DNA-DDA is a nonlinear dynamics method for predicting A/B compartments based on the DNA sequence alone. We derived DNA-DDA models for four human cell types using only a 20 Mb-long region on chr22 and corresponding compartment labels defined by Hi-C data. DNA-DDA achieved a mean AUCs across all held-out chromosomes between 0.75 and 0.84. The achieved performances on never-before-seen testing data demonstrate the potential of DDA, a method which has been shown to accurately classify time series data, in the field of genomics.

Many computational methods for predicting genome architecture have been developed in recent years [62], but only a few are able to predict A/B compartments from the sequence alone. ABCNet [59] and SACSANN [8] are both convolutional neural network (CNN) based models that can predict A/B compartments. While ABCNet relies only on the sequence, SACSANN [8] uses counts of sequence-derived features (e.g., TEs, TFBSs). Both achieve AUC scores close to 80%, but while ABCNet does not require any previous knowledge about the genome of interest, SACSANN derives its input features from annotation data sets that are not available for every species. The currently the most comprehensive sequence-based approach for modeling 3D genome architecture is Orca [60]. Orca is also the first sequence based model able to predict truly long-range interactions (> 1 Mb). Orca models have been trained on two of the most highest resolution microC data sets to date, and predict interactions simultaneously at different resolutions. The models’ ability to capture sequence dependencies of 3D genome architecture has been experimentally validated. DNA-DDA competes well with these state-of-the-art methods which use a larger portion of the genome for training and/or input features other than solely the sequence (Table 4).

This study is a proof of concept meant to illustrate the potential in applying DDA-based approaches in the field of structural genomics. We want to emphasize that the DNA-DDA models presented here highly unlikely represent an optimum. Many aspects of our analysis could be evaluated and modified, including the use of a different numerical DNA sequence representation, other functional forms for the DDA models, and alternative targets to assess model performance.

We encoded the DNA sequence as a 1D DNA walk [54, 63], which is a common representation for analysis of DNA sequences in time series frameworks and has been the basis for numerous studies that applied spectral analysis and signal processing methods such as discrete or Ramanujan fourier transform and wavelet and fractal analysis for revealing high-level periodicities and patterns with biological significance [33, 34, 64, 65]. DDA analysis can be related to spectral analysis: the estimated features and time delays of linear DDA models relate directly to frequencies in the data, and in nonlinear DDA models, they relate to higher order statistical moments [36].

In the 1D DNA walk using the *hydrogen bond energy* (*SW*) *rule* [54, 66], the walker starts at zero and continues along the linear chain of nucleotides taking a step up for strongly bonded pairs (C or G) and down for weakly bonded pairs (A or T). Thus, DNA-DDA models capture dynamical properties based mainly on GC-content. Of course, more complex representations of the DNA sequence have been proposed as well, some of which take all four nucleotides into account (overview and comparison in [64, 67, 68]). We initially considered the alternative and equally simple 1D mapping integer representation (*T* = 0, *C* = 1, *A* = 2, *G* = 3). However this method implies biologically irrelevant properties on the bases such that purines are weighted more than pyrimidines ((*A,G*) > (*C,T*)). In future work, we plan to resort to DNA representations that include information of all nucleotides and do not have such a bias such as the 2D DNA walk [69]. Naturally, the ergodicity measure has to be substantially modified to achieve this.

The functional form of the DNA-DDA model was chosen based only on simplicity and computational efficiency. Previous work has shown that the overall functional form of a DDA model tends to be specific to the data type used to measure the system of interest (e.g., EEG, ECG, DNA sequence), while the delay pairs are sensitive to the question we ask about the system [58]. A large-scale exhaustive sweep of model-delay-pair combinations such as described in Lainscsek *et al* [41] could be implemented to optimize model-delay pair combinations.

To predict A/B compartments, DNA-DDA first constructs a (DNA-DDA) contact matrix. This matrix is then postprocessed (e.g., filtering, logarithmic transformation, etc.) before being subjected to PCA, as proposed by Lieberman *et al* [1]. The Although compartments are routinely identified using PCA, the binary classification into A and B compartments is most likely an oversimplification of the 3D genome architecture [5]. In fact, Rao *et al* [2] showed that genomes segregate into at least six subcompartments, each exhibiting a distinctive pattern of genomic and epigenetic features. Furthermore, we chose the PC to be most representative of compartments based on its correlation with a histone mark associated with open chromatin in the respective cell type and genomic region of interest, as suggested by [4]. However, the chosen PC is clearly dependent on the region being considered and often, two or more PCs will often exhibit very similar correlation coefficients to the ChIP-seq signal. Naturally, this is not a limitation exclusive to DNA-DDA, but rather, of all methods calling compartments based on PCA. A large improvement in the structure selection step would be to select delay-pairs based on an alternative mutli-variate compartment classification method or the similarity of the DNA-DDA and Hi-C contact matrices directly. Since all relevant 3D structures could be extracted from interaction matrices (at various resolutions), we strongly believe that choosing the delay-pairs with a better suited target label will substantially boost cell-type specificity.

DDA models are sparse and comprise a small number of features (typically 1-4), making them robust to overfitting and well generalizable to new data [36]. Large models such as deep learning networks run the risk of capturing irrelevant patterns or “noise” in the data. In contrast, small and simple models typically fail to capture dominant signatures in the data. Since DDA does not look for the most predictive, but rather the most discriminative model, most terms in Equation 1 are set to zero. This is the power of DDA, it does not aim to model but rather capture dominant dynamical signatures in the data and 3-term DDA models have been proven sufficient for classifying complex biological data sets (eg. [41, 58, 70]). Still, one caveat of DNA-DDA concerns feature interpretability. Although models with fewer parameters are often more interpretable than the immense feature spaces that are typical of deep learning, this is not the case with DDA due to its foundation and motivation in embedding theory [37]. The estimated coefficients and delays of a linear DDA model relate to the frequencies of a signal, and the estimated coefficients of nonlinear DDA models are connected to higher-order statistical moments [36]. In general, the delays of nonlinear DDA models are characterized by complex phase and frequency couplings which are not attributable to one particular system property. Nevertheless, due to the extremely low computational load of DNA-DDA, sequencebased mechanisms and genetic variants, such mutations in known or putative binding sites for regulatory proteins or genomic rearrangements, could easily be tested. Importantly, a similar issue arises concerning the interpretability of deep learning based models. It would be interesting to see what information about 3D genome architecture could be recovered with a combined effort of deep learning-based and nonlinear dynamics-based methodologies.

The power of methods like DNA-DDA and Orca lies in their intermediate step of predicting the contact matrix before calling A/B compartments, since all relevant 3D structures could be extracted from them in the same manner as Hi-C maps. Although Orca currently predicts contact maps for higher resolutions, DNA-DDA shows promising results whilst tackling the problem from a different angle and requiring far less “training data”. We chose a relatively low resolution (100 kb) for this proof-of-concept study to 1) efficiently test the many possible parameters and aspects of our approach; and 2) allow for a better comparison with SACSANN and ABCNet, which both operate at 100 kb, and together with Orca are the only other purely sequence based methods to predict compartments of which we are aware. We acknowledge that our current DNA-DDA contact maps capture global large-scale patterns rather than subtle cell-type specific changes. Nevertheless, we believe that the use of a different numerical DNA sequence representation and/or other DDA model forms, and alternative targets to assess model performance will greatly improve celltype specificity and result in more diverse DNA-DDA contact maps (Figure 4). Moreover, our results show that DDA models can be derived from a small amount of supervised data, which would enable the prediction of genome-wide interactions in other cell types from a very limited amount of chromosome conformation capturing data (e.g., generated with 5C). With some optimization, we are convinced that the method might be able to do this as well as deep learning algorithms, but using just a miniscule fraction of the volume of data required by such approaches.

**Fig. 3.**
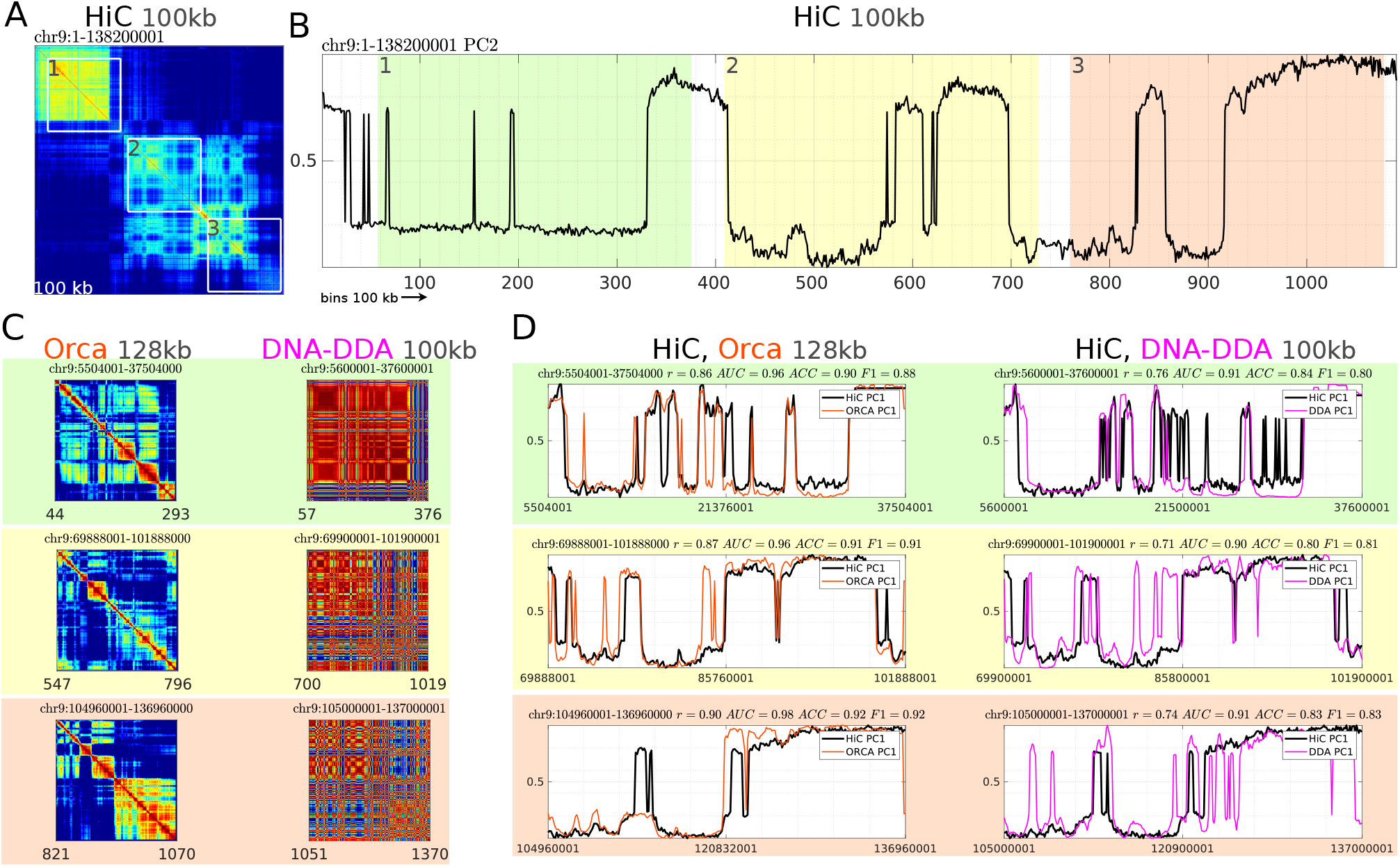
Comparison of DNA-DDA and Orca for the hESC cell line. A) hESC Hi-C contact map of chr9 at 100 kb resolution and three highlighted regions (1, 2, and 3) which were used for comparison. The PC with the highest correlation to the H3K4me1 is PC2. B) PCA on the entire Hi-C contact map in A). C) Orca and DNA-DDA Pearson correlation matrices for regions 1, 2 and 3. The regions were slightly increased to accommodate the Orca model, which predicts contacts at 128 kb resolution. D) PCA analysis on regions 1, 2 and 3 individually for 128 kb (left) and 100 kb (right) resolution for Hi-C data (black), Orca (orange), and DNA-DDA (magenta). In all cases and for all regions, the PC with the highest correlation to H3K4me1 was PC1. The correlation coefficients for different PCs (here, PC1 and PC2) are often very similar.

**Fig. 4.**
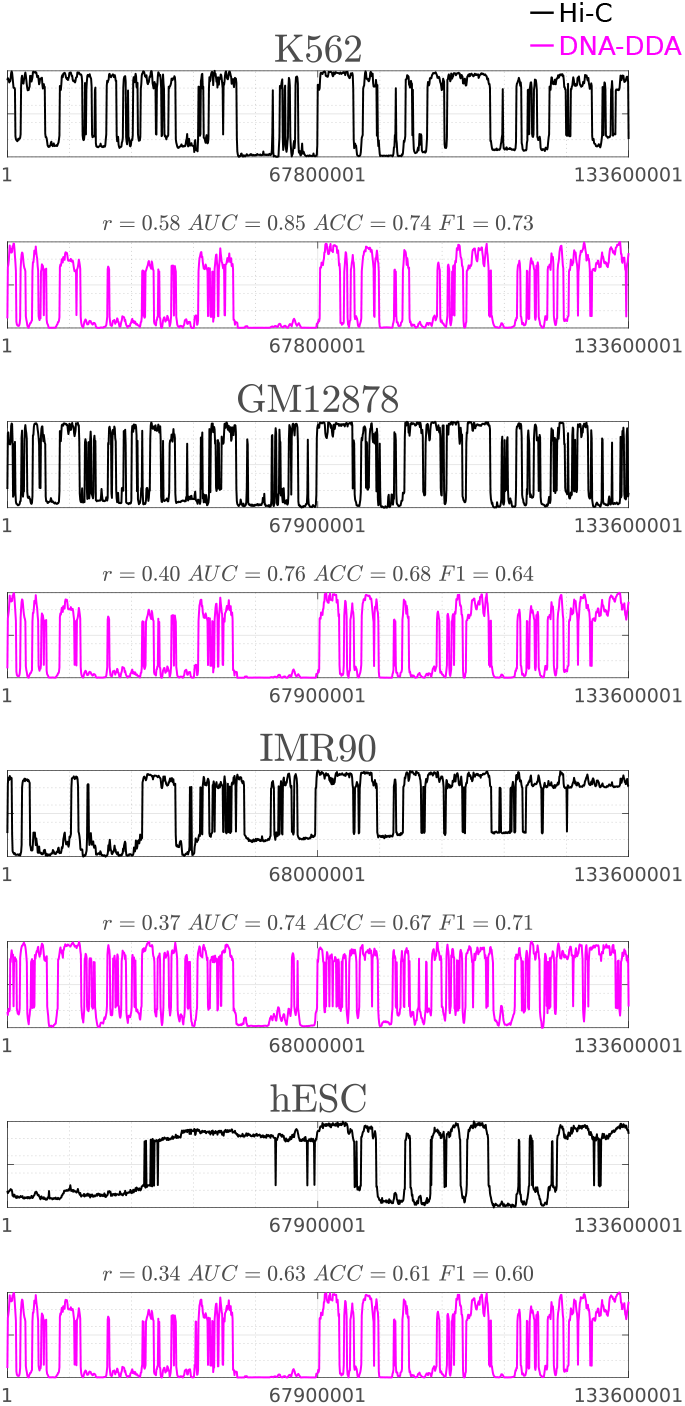
Comparison of DNA-DDA models in for four cell types. Principal components PC_DNA-DDA_ (magenta) and PC_Hi-C_ (black) are shown for chr10 of each cell line (K562, GM12878, IMR90, hESC). The overall Pearson correlation coefficient between the PC_Hi-C^s^_ across all cell lines is 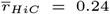 where GM12878 and K562 are the most similar 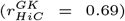, and K562 and hESC are the most different 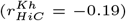. The overall Pearson correlation coefficient between the PC_DNA-DDA^s^_ across all cell lines is 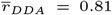 where GM12878 and hESC are the most similar 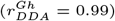, and IMR90 and GM12878 are the most different 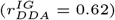.

The DNA sequence plays a fundamental role in the formation and maintenance of 3D genome architecture, which in turn is a central orchestrator of the gene regulatory network. It is indisputable that the sequence contains key underlying features contributing to genome folding, however to what extent it alone can be used to gain access to all 3D interactions and relevant structures remains an open question. Our findings strongly support the hypothesis that the DNA sequence represents a highly observable variable of chromatin architecture and that chromosomal compartments can be predicted solely from the DNA sequence. This opens up a variety of possibilities such as discovering novel sequence signatures imperative to structural genome function and how disruption of 3D genome architecture relates to human disease.

### Key Points

- Substantial information about 3D genome architecture can be uncovered from from solely the DNA sequence.
- DNA-DDA models derived from fraction of the typical amount of labeled data needed, already show high predictive power in classifying A/B compartments.
- Delay differential analysis, a technique with foundations in chaos theory, is suited for analyzing genomic sequence data in the context of structural genomics.

## Acknowledgements

We thank Dr. Claudia Lainscsek for the fruitful discussions and sharing the C implementation of DDA, which was used as a basis for DNA-DDA.

## Data availability

Exemplary code used to predict contacts at 100 kb resolution of chr22 can be found at https://github.com/xX3N1A/DNA-DDA.

## References

1. Lieberman-Aiden E, van Berkum N L, Williams L, et al. Comprehensive mapping of long-range interactions reveals folding principles of the human genome. Science. 2009;326(5950):289–93.

2. Rao SS, Huntley MH, Durand NC, et al. A 3D map of the human genome at kilobase resolution reveals principles of chromatin looping. Cell. 2014;159(7):1665–80.

3. Liu Y, Nanni L, Sungalee S, et al. Systematic inference and comparison of multi-scale chromatin sub-compartments connects spatial organization to cell phenotype. Nature Communications. 2021;12:2439.

4. Fortin J, Hansen K. Reconstructing A/B compartments as revealed by Hi-C using long-range correlations in epigenetic data. Genome Biology. 2015;16(180).

5. Nichols MH, Corces VG. Principles of 3D compartmentalization of the human genome. Cell Reports. 2021;35(13):109330.

6. Corbo M, Damas J, Bursell MG, et al. Conservation of chromatin conformation in carnivores. PNAS. 2022;119:e2120555119.

7. Feurtey A, Lorrain C, Croll D, et al. Genome compartmentalization predates species divergence in the plant pathogen genus Zymoseptoria. BMC Genomics. 2020;21:588.

8. Prost J, Cameron C, Blanchette M. SACSANN: identifying sequence-based determinants of chromosomal compartments. bioRxiv. 2020; doi: 10.1101/2020.10.06.328039.

9. Krijger PHL, de Laat W. Regulation of disease-associated gene expression in the 3D genome. Nature reviews Molecular cell biology. 2016;17(12):771–782.

10. Gorkin DU, Qiu Y, Hu M, et al. Common DNA sequence variation influences 3-dimensional conformation of the human genome. Nature reviews Molecular cell biology. 2019;20(1):255.

11. Krumm A, Duan Z. Understanding the 3D genome: Emerging impacts on human disease. Seminars in cell & developmental biology. 2019;90:62–77.

12. Degn H, Holden AV, Olsen LF. Chaos in Biological Systems. In: Degn H, Holden AV, Olsen LF (eds). NATO Advanced Research Workshop on “Chaos in Biological Systems” December 8-12, 1986. Dyffryn House, St. Nicholas, Cardiff, U. K., 1987, 1–171.

13. Letellier C. Chaos in Nature 2nd Edition. Singapore: WORLD SCIENTIFIC, 2013.

14. Hewelt B, Li H, Jolly MK, et al. The DNA walk and its demonstration of deterministic chaos-relevance to genomic alterations in lung cancer. Bioinformatics. 2019;35(16):2738–2748.

15. Lorenz E. Deterministic Nonperiodic Flow. Journal of Atmospheric Sciences. 1963;20(2):130–148.

16. Dias F, Iooss G. Chapter 10 - Water-Waves as a Spatial Dynamical System. In: Friedlander S, Serre D (eds). Handbook of Mathematical Fluid Dynamics. vol. 2 of Handbook of Mathematical Fluid Dynamics. North-Holland, 2003, 443–499.

17. Morfu S, Marquié P, Nofiele B, et al. Nonlinear Systems for Image Processing. Advances in Imaging and Electron Physics. 2008 01; doi: 10.1016/S1076-5670(08)00603-4.

18. Poincaré H. Sur le problème des trois corps et les équations de la dynamique. Acta Math. 1890;13:1–270.

19. Poincaré H. Methodes nouvelles de la mécanique celeste. Paris: Gauthier-Villars et fils, 1854-1912.

20. Ruelle D. Strange attractors. The Mathematical Intelligencer. 1980;2:126–137.

21. Grebogi C, Ott E, Pelikan S, et al. Strange attractors that are not chaotic. Physica D: Nonlinear Phenomena. 1984;13(1):261–268.

22. Lefranc M. The Topology of Deterministic Chaos: Stretching, Squeezing and Linking. Physics and Theoretical Computer Science. 2007 01;7:71–90.

23. Lyapunov AM. The General Problem of the Stability of Motion. PhD thesis. University of Moscow, 1892.

24. Mandelbrot BB. Les objets fractals: forme, hasard et dimension. Paris: Flammarion, 1975.

25. Boltzmann L. Vorlesungen uber Gastheorie. Bd. 2. Leipzig: Verlag von Johann Ambrosius Barth, 1889.

26. Birkhoff GD. Proof of the Ergodic Theorem. Proceedings of the National Academy of Sciences. 1931;17(12):656–660.

27. v Neumann J. Proof of the Quasi-Ergodic Hypothesis. Proceedings of the National Academy of Sciences. 1932;18(1):70–82.

28. Shields PC. String Matching: The Ergodic Case. The Annals of Probability. 1992;20(3):1199–1203.

29. Falconnet M, Gantert N, Saada E. Ergodicity of some dynamics of DNA sequences. arXiv. 2019; doi: missing?!?

30. Shannon CE. An algebra for theoretical genetics. PhD thesis. Massachusetts Institute of Technology - Department of Mathematics, 1940.

31. Chanda P, Costa E, Hu J, et al. Information Theory in Computational Biology: Where We Stand Today. Entropy. 2020;22(6).

32. Lobzin VV, Chechetkin VR. Order and correlations in genomic DNA sequences. The spectral approach. Physics-Uspekhi. 2000;43:55–78.

33. Weighill D, Macaya-Sanz D, DiFazio SP, et al. Wavelet-Based Genomic Signal Processing for Centromere Identification and Hypothesis Generation. Frontiers in genetics. 2019;10:487.

34. Yin C, Chen Y, Yau STS. A measure of DNA sequence similarity by Fourier Transform with applications on hierarchical clustering. Journal of theoretical biology. 2014;359:18–28.

35. S V. Information theory applications for biological sequence analysis. Briefings in Bioinformatics. 2014;15:376–389.

36. Lainscsek C, Sejnowski TJ. Delay Differential Analysis of Time Series. Neural Computation. 2014;23(3):594–614.

37. Takens F. Detecting strange attractors in turbulence. In: Rand D, Young LS (eds). Dynamical Systems and Turbulence, Warwick 1980. Berlin, Heidelberg: Springer Berlin Heidelberg, 1981, 366–381.

38. Aguirre I, Letellier C. Investigating observability properties from data in nonlinear dynamics. Physical review E, Statistical, nonlinear, and soft matter physics. 2020;83.

39. Lainscsek C, Cash SS, Sejnowski TJ, et al. Dynamical ergodicity DDA reveals causal structure in time series. Chaos. 2021;31:103108.

40. Eckmann JP, Oliffson Kamphorst S, Ruelle D. Recurrence Plots of Dynamical Systems. Europhysics Letters (EPL). 1987;4(9):973–977.

41. Lainscsek C, Weyhenmeyer J, Cash SS, et al. Delay Differential Analysis of Seizures in Multichannel Electrocorticography Data. Neural Computation. 2017;29(12):3181–3218.

42. Lainscsek C, Gonzalez CE, Sampson AL, et al. Causality detection in cortical seizure dynamics using cross-dynamical delay differential analysis. Chaos. 2019;29:101103.

43. Lainscsek C, Rungratsameetaweemana R, Cash SS, et al. Cortical chimera states predict epileptic seizures. Chaos. 2019;29:121106.

44. Sanborn AL, Rao SS, Huang SC, et al. Chromatin extrusion explains key features of loop and domain formation in wildtype and engineered genomes. Proc Natl Acad Sci U S A. 2015;112(47):E6456–65.

45. ENCODE Project Consortium. An integrated encyclopedia of DNA elements in the human genome. Nature. 2012;489(7414):57–74.

46. Karolchik D, Hinrichs AS, Furey TS, et al. The UCSC Table Browser data retrieval tool. Nucleic Acids Res. 2004;1(32):D493–6.

47. Dixon JR, Selvaraj S, Yue F, et al. Topological domains in mammalian genomes identified by analysis of chromatin interactions. Nature. 2012;485(7398):376–80.

48. Parry AJ, Hoare M, Bihary D, et al. NOTCH-mediated non-cell autonomous regulation of chromatin structure during senescence. Nat Commun. 2018;9(1):1840.

49. Ben L, Salzberg SL. Fast gapped-read alignment with Bowtie 2. Nature Methods. 2012;9:357–359.

50. Wolff J, Rabbani L, Gilsbach R, et al. Galaxy HiCExplorer 5: a web server for reproducible Hi-C, capture Hi-C and single-cell Hi-C data analysis, quality control and visualization. Nucleic Acids Research. 2020;48(W1):W177–W184.

51. Wolff J, Bhardwaj V, Nothjunge S, et al. Galaxy HiCExplorer: a web server for reproducible Hi-C data analysis, quality control and visualization. Nucleic Acids Research. 2018;46(W1):W11–W16.

52. Ramirez F, Bhardwaj V, Arrigoni L, et al. High-resolution TADs reveal DNA sequences underlying genome organization in flies. Nature Communications. 2018;9:189.

53. Knight AP, Ruiz D. A Fast Algorithm for Matrix Balancing. IMA J Numer Anal. 2007;33.

54. Buldyrev SV, Goldberger AL, Havlin S, et al. Long-range correlation properties of coding and noncoding DNA sequences: GenBank analysis. Physical review E, Statistical physics, plasmas, fluids, and related interdisciplinary topics. 1995;51(5):5084–5091.

55. Heinz S, Benner C, Spann N, et al. Simple combinations of lineage-determining transcription factors prime cis-regulatory elements required for macrophage and B cell identities. Mol Cell. 2010;38:576–89.

56. Tsompana M, Buck MJ. Chromatin accessibility: a window into the genome. Epigenetics & Chromatin. 2014;7(33).

57. Lainscsek C, Hernandez ME, Weyhenmeyer J, et al. Nonlinear dynamical analysis of EEG time series distinguishes patients with Parkinson’s disease from healthy individuals. Frontiers in neurology. 2013;4.

58. Sampson AL, Lainscsek C, Gonzalez CE, et al. Delay differential analysis for dynamical sleep spindle detection. Journal of neuroscience methods. 2019;316:12–21.

59. Kirchhof M, Cameron CJ, Kremer SC. End-to-end chromosomal compartment prediction from reference genomes. In: 2021 IEEE International Conference on Bioinformatics and Biomedicine (BIBM), 2021, 50–57.

60. Zhou J. Sequence-based modeling of three-dimensional genome architecture from kilobase to chromosome scale. Nat Genetics. 2022;54:725–734.

61. Krietenstein N, Abraham S, Venev S, et al. Ultrastructural Details of Mammalian Chromosome Architecture. Mol Cell. 2020;78(3):554–565.

62. Belokopytova P, Fishman V. Predicting Genome Architecture: Challenges and Solutions. Frontiers in genetics. 2021;11:617202.

63. Peng CK, Buldyrev SV, Goldberger AL, et al. Long-range correlations in nucleotide sequences. Nature. 1992;356:168–170.

64. Mendizabal-Ruiz G, Román-Godmez I, Torres-Ramos S, et al. On DNA numerical representations for genomic similarity computation. PLOS ONE. 2017;12(3):1–27.

65. Haimovich AD, Byrne B, Ramaswamy R, et al. Wavelet analysis of DNA walks. Journal of computational biology. 2006;13(7):1289–1298.

66. Berger JA, Mitra SK, Carli M, et al. Visualization and analysis of DNA sequences using DNA walks. Journal of the Franklin Institute. 2004;341(1):37–53.

67. Kwan HK, Arniker SB. Numerical representation of DNA sequences. In: IEEE International Conference on Electro-Information Technology, 2009, 307–310.

68. Kumar GS. DNA Sequence Representation methods. In: ISB Proceedings of the International Symposium on Biocomputing, 2009, 1–4.

69. Zhang L, Jiang Z. Long-range correlations in DNA sequences using 2D DNA walk based on pairs of sequential nucleotides. Chaos, Solitons & Fractals. 2004;22(4):947–955.

70. Lainscsek C, Weyhenmeyer J, Hernandez ME, et al. Nonlinear dynamical classification of short time series of the rossler system in high noise regimes. Frontiers in neurology. 2013;4:182.

