## Supplementary Material for "Predicting chromosomal compartments directly from the nucleotide sequence with DNA-DDA"

Xenia Lainscsek [1] and Leila Taher [1]

[1]Institute of Biomedical Informatics, Graz University of Technology

### Figures

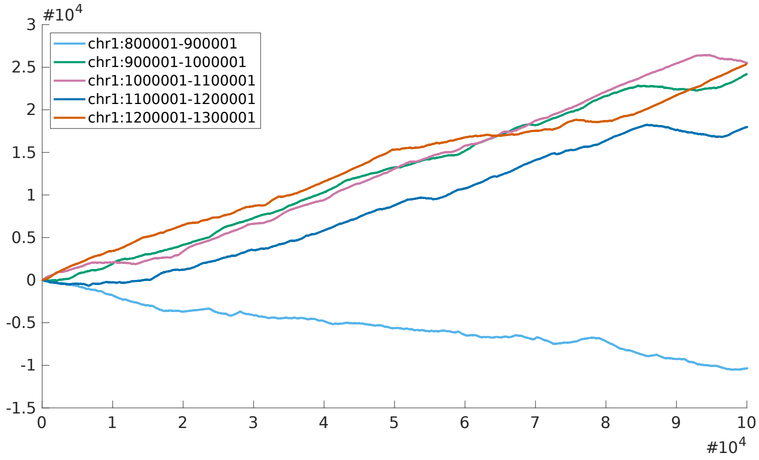


Figure S1: Random walk representation for first five bins of chr1

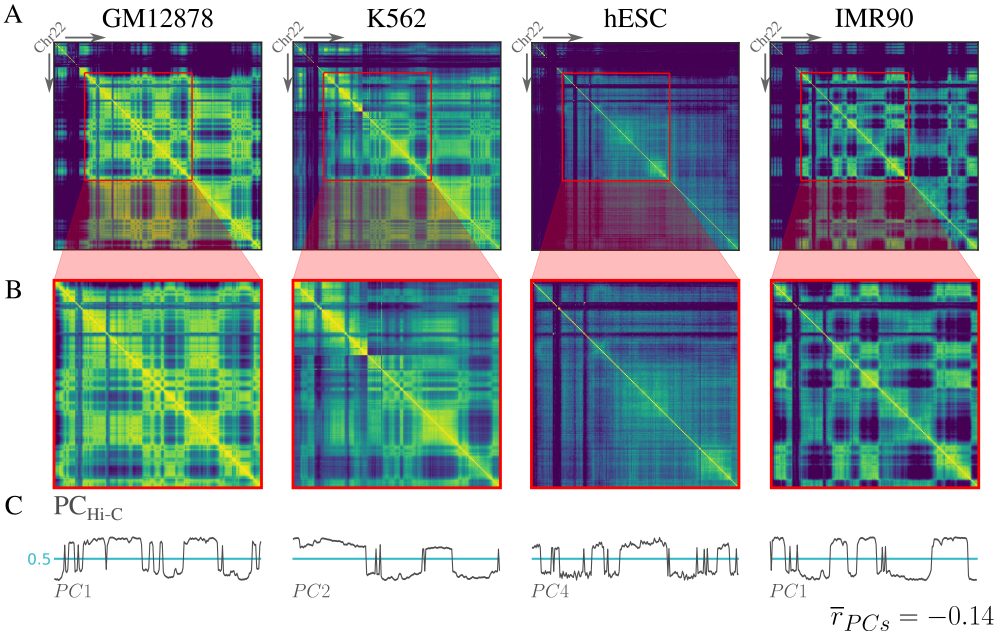


Figure S2: **Data used for supervised structure selection** A) The reference sequence of human chromosome 22 has 50,818,467 bps and can be partition into 509 100kbp-long bins. B) The 200 consecutive bins (chr22:16200000:36200001) with the C) highest variation (lowest Pearson's $\overline{r}_{low} = -0.14$) between cell types were selected to be used for structure selection. The PC number chosen to represent chromosomal compartments is based on their correlation with the H3K4me profile in the respective cell line. This can change depending on which region is being considered. Column two in the table indicates $PC_{chr22:1-50818468} \rightarrow PC_{chr22:16200000:36200001}$: K562 $PC_1 \rightarrow PC_2$; hESC $PC_1 \rightarrow PC_4$

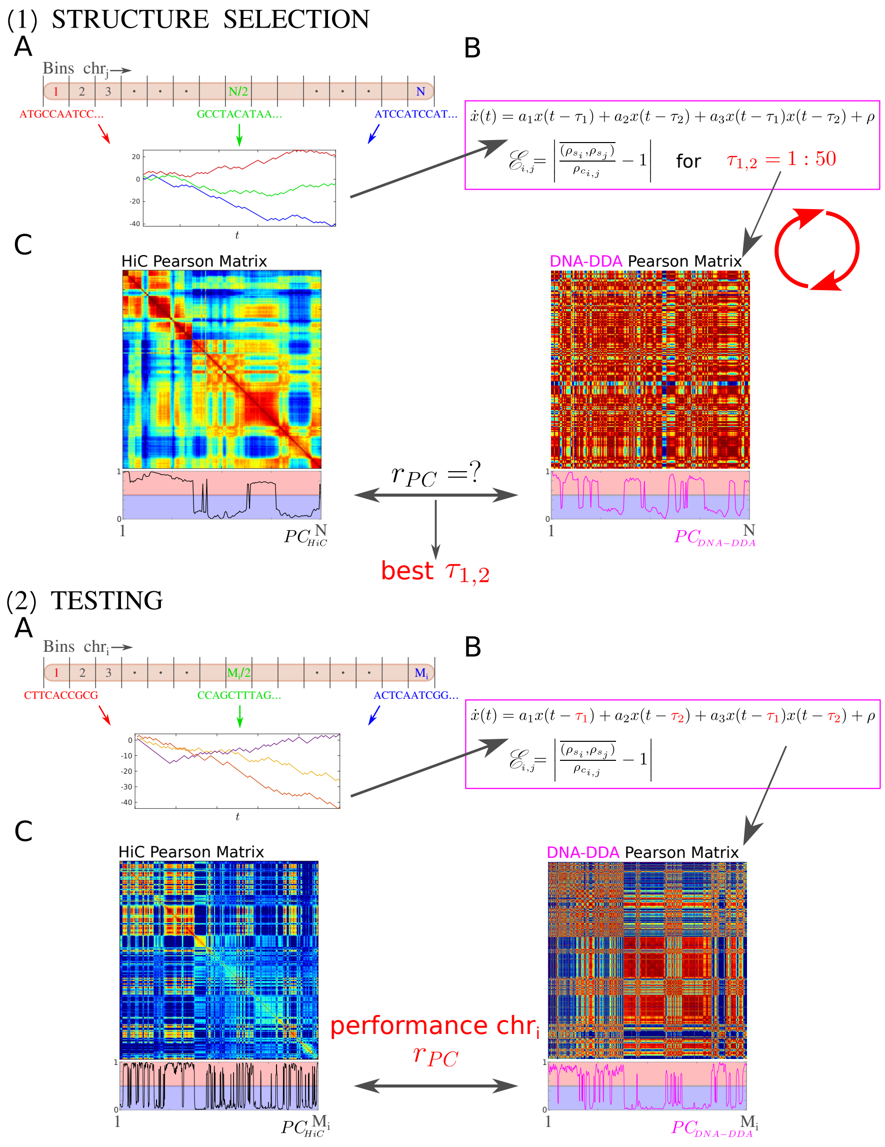


Figure S3: **Workflow for determining DNA-DDA parameters $\tau_1$ and $\tau_2$ and subsequent testing for an arbitrary cell line. 1) Structure Selection** 1A) Numerical encoding of the sequence as a 1D DNA walk $x(t)$ using structure selection data on chromosome $j$: $chr_j : 1 - N$ 1B) $x(t)$ is inputted into the DNA-DDA model; the ST and CT errors $\rho_{ST}, \rho_{CT}$ are computed and combined to DE-DDA $\mathscr{E}$ (see Figure 1 ) for all delay pair combinations $\tau_1, \tau_2 \in [1 : 50]$ 1C) the best performing delay pair, defined as that which gives the highest Pearson correlation coefficient $r_{PC}$ between $PC_{\text{Hi-C}}$ and $PC_{\text{DNA-DDA}}$, is fixed. **2) Testing** 2A) Numerical encoding of the sequence as a 1D DNA walk $x(t)$ for an arbitrary hold-out testing chromosome $i$: $chr_i : 1 - M$ 2B) $x(t)$ is inputted into the DNA-DDA model; the ST and CT errors $\rho_{ST}, \rho_{CT}$ are computed and combined to DE-DDA $\mathscr{E}$ for best performing delay pair $\tau_1, \tau_2$ determined in step 1 2C) the final performance for DNA-DDA on chromosome $i$ is given by $r_{PC}$ between $PC_{\text{Hi-C}}$ and $PC_{\text{DNA-DDA}}$.

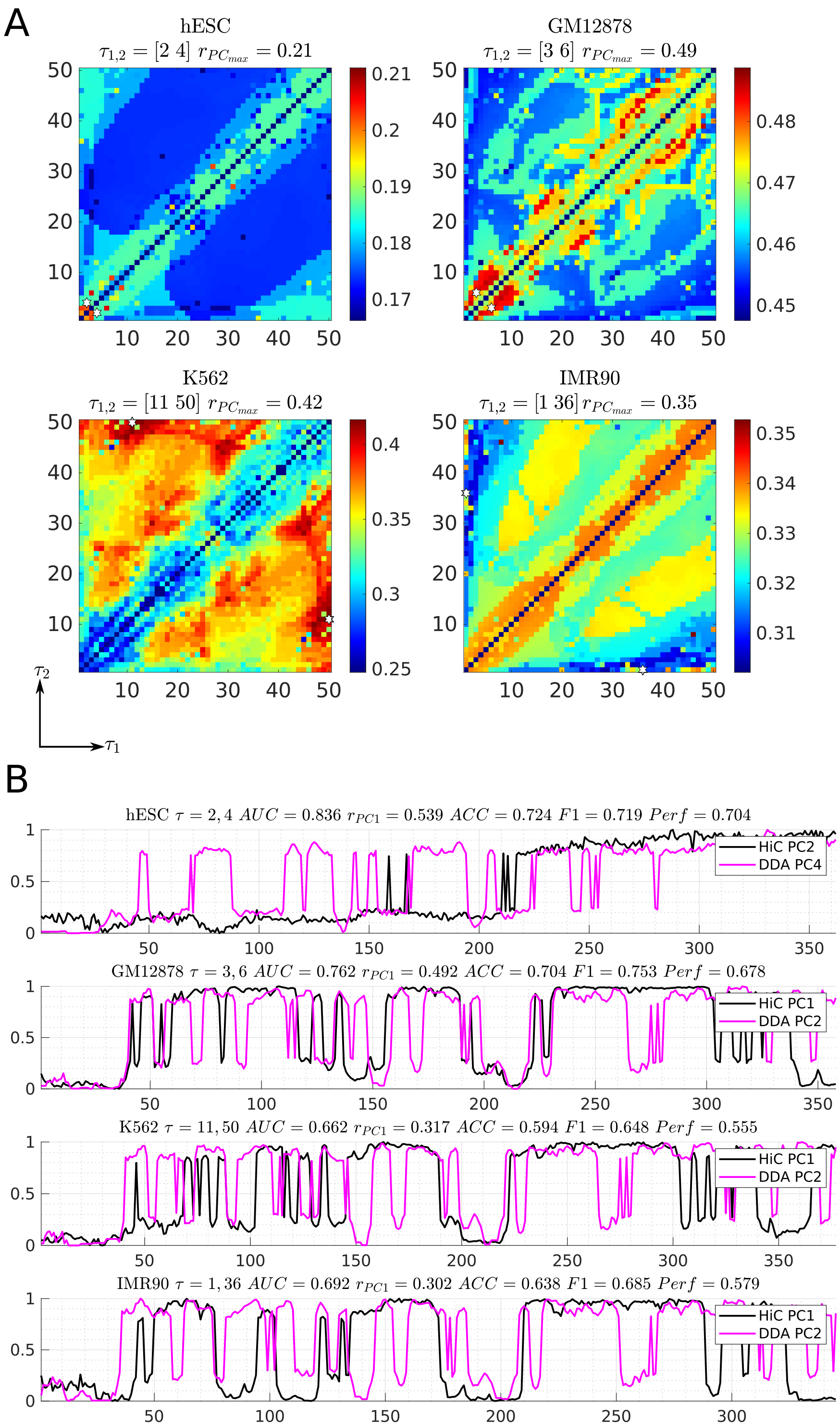


Figure S4: **Supervised structure selection on chr22 reveals regions of enriched dynamics across delay pairs and cell types.** A) Heatmaps depicting performance (Pearson correlation between $PC_{\text{DNA-DDA}}$ and $PC_{\text{Hi-C}}$) for each delay pair $\tau_1, \tau_2$ between 1 and 50 on genomic region chr22:16200000:36200001. The best performing delay pairs are indicated by a white hexagram. B) Performance of best performing delay pairs over entire chromosome 22.

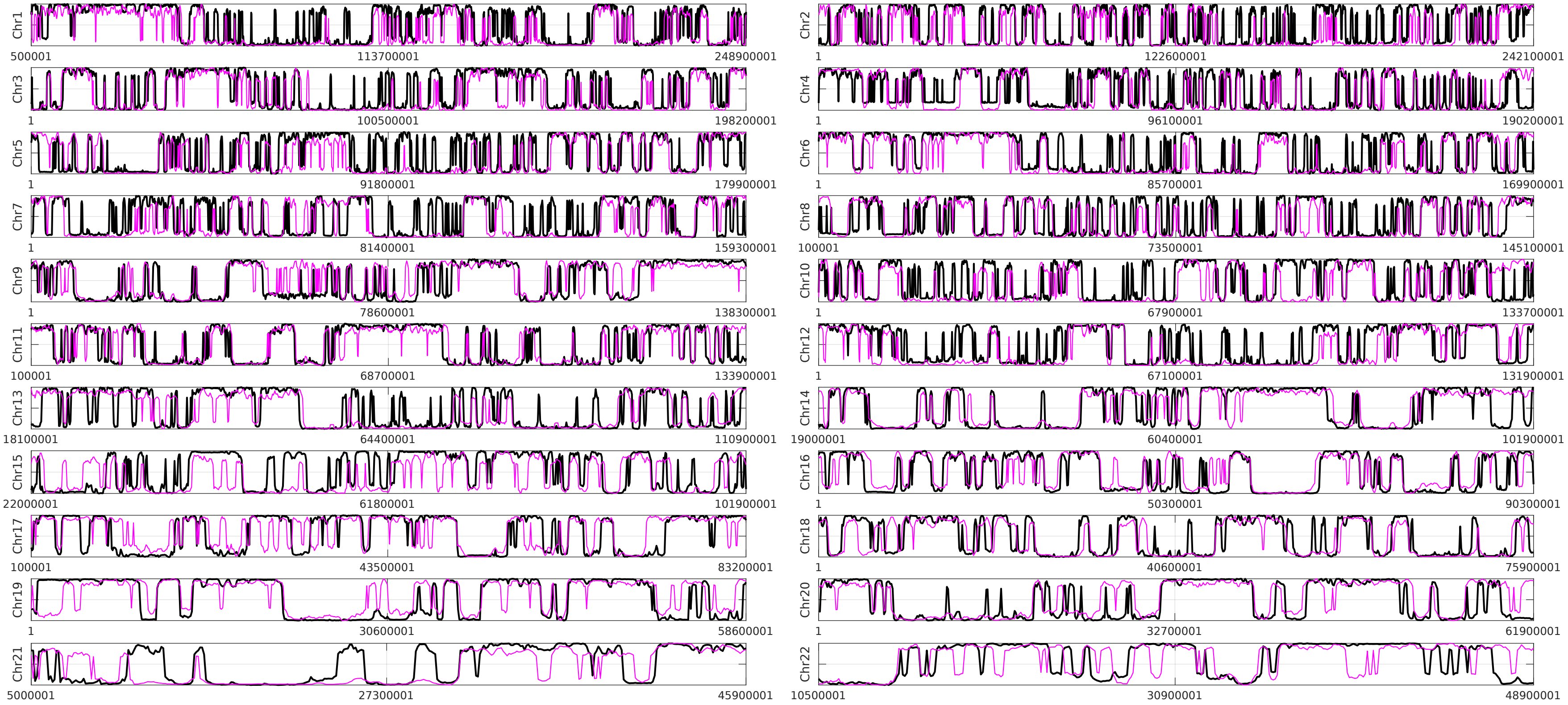


Figure S5: **$\text{PC}_{\text{DNA-DDA}}$ (magenta) and $\text{PC}_{\text{Hi-C}}$ (black) for GM12878 over each chromosome. normalized to 0 and 1.** Values above 0.5 correspond to the A compartment; values below 0.5 correspond to the B compartment.

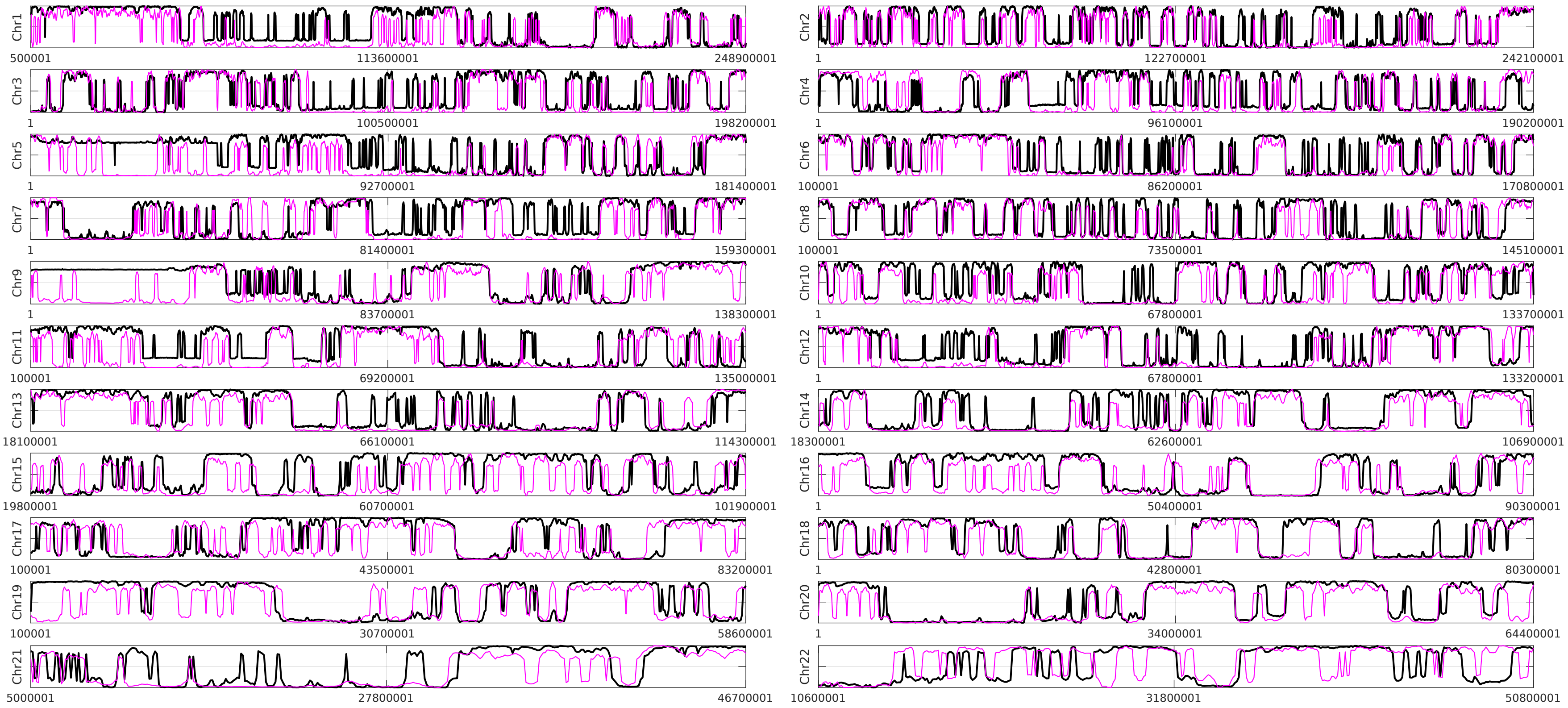


Figure S6: **$PC_{DNA\text{-}DDA}$ (magenta) and $PC_{Hi\text{-}C}$ (black) for K562 over each chromosome. normalized to 0 and 1.** Values above 0.5 correspond to the A compartment; values below 0.5 correspond to the B compartment.

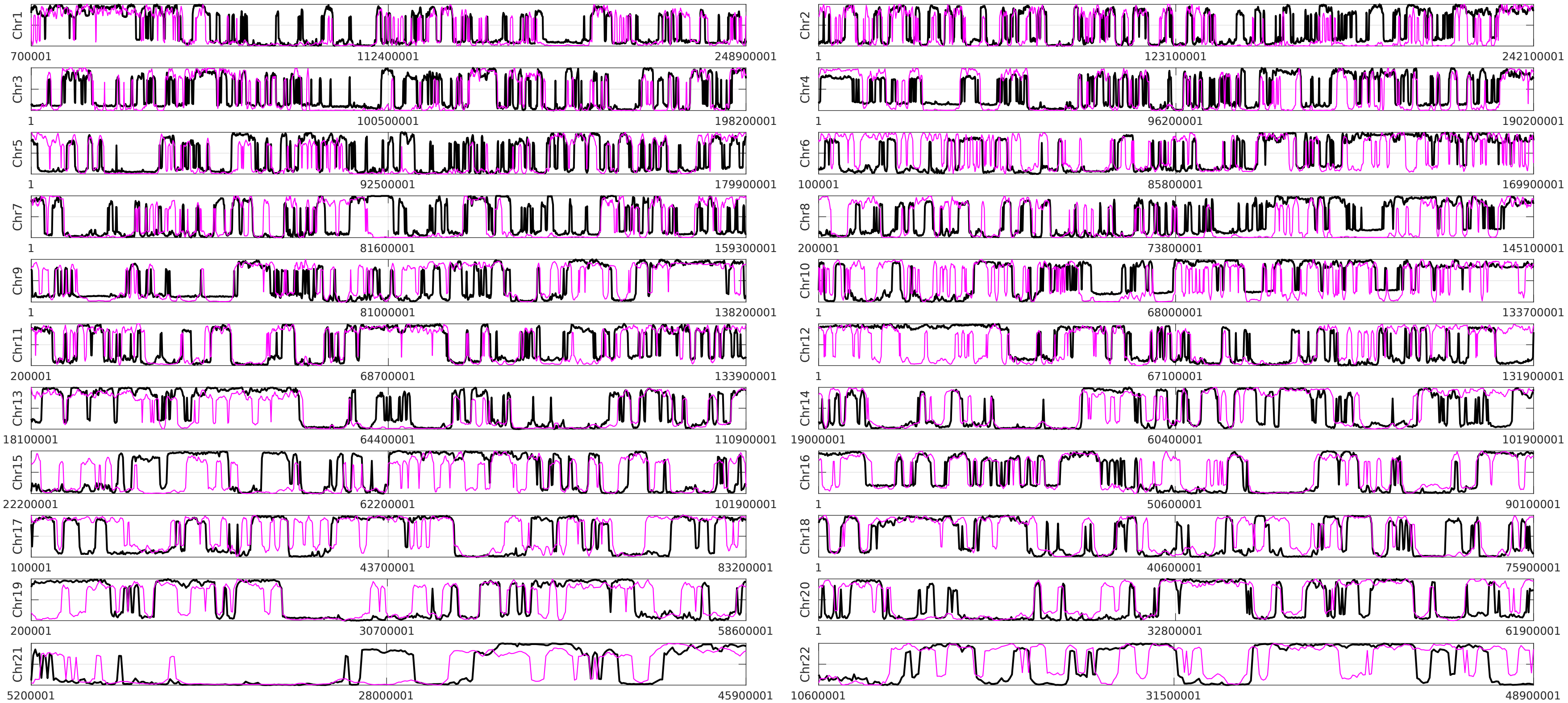


Figure S7: **$PC_{\text{DNA-DDA}}$ (magenta) and $PC_{\text{Hi-C}}$ (black) for IMR90 over each chromosome normalized to 0 and 1.** Values above 0.5 correspond to the A compartment; values below 0.5 correspond to the B compartment.

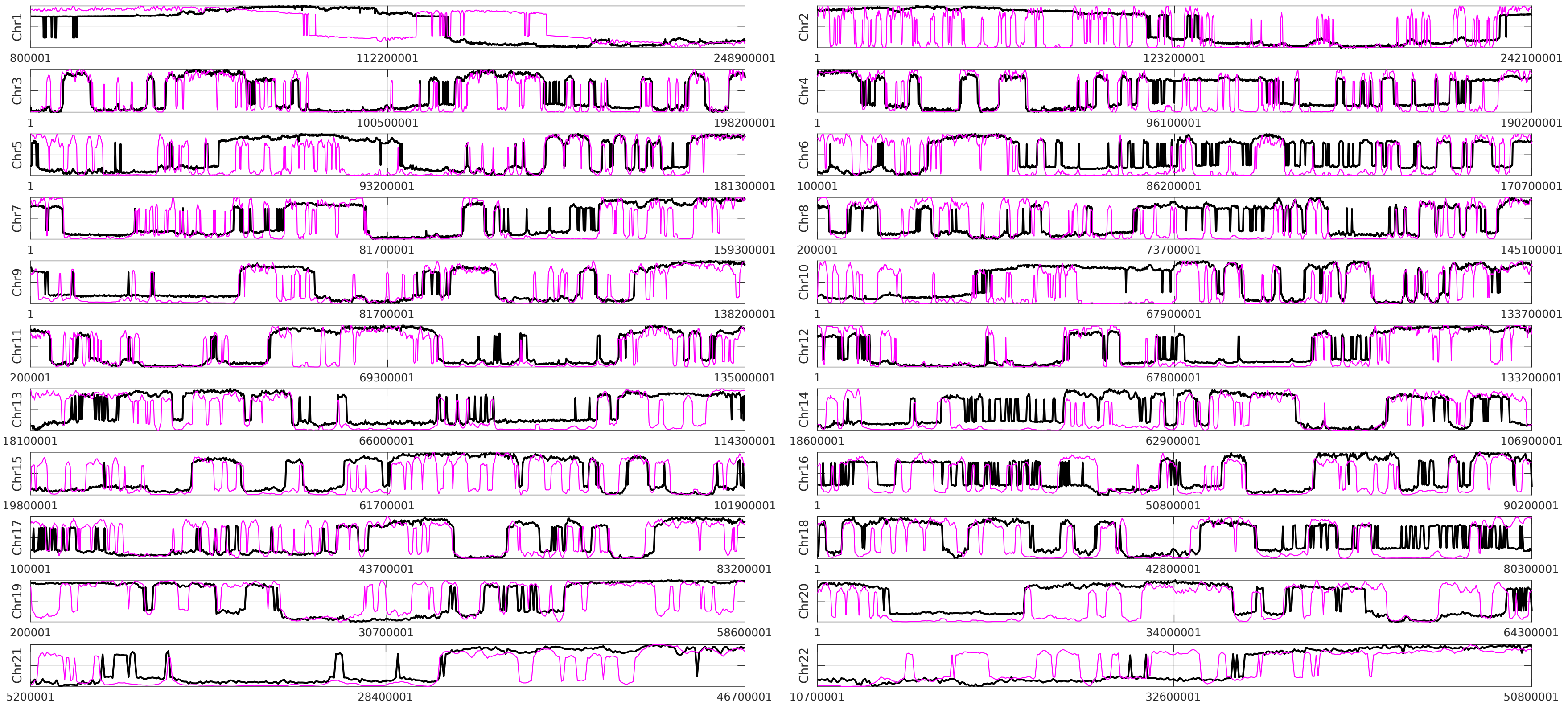


Figure S8: **$\text{PC}_{\text{DNA-DDA}}$ (magenta) and $\text{PC}_{\text{Hi-C}}$ (black) for hESC over each chromosome normalized to 0 and 1.** Values above 0.5 correspond to the A compartment; values below 0.5 correspond to the B compartment.

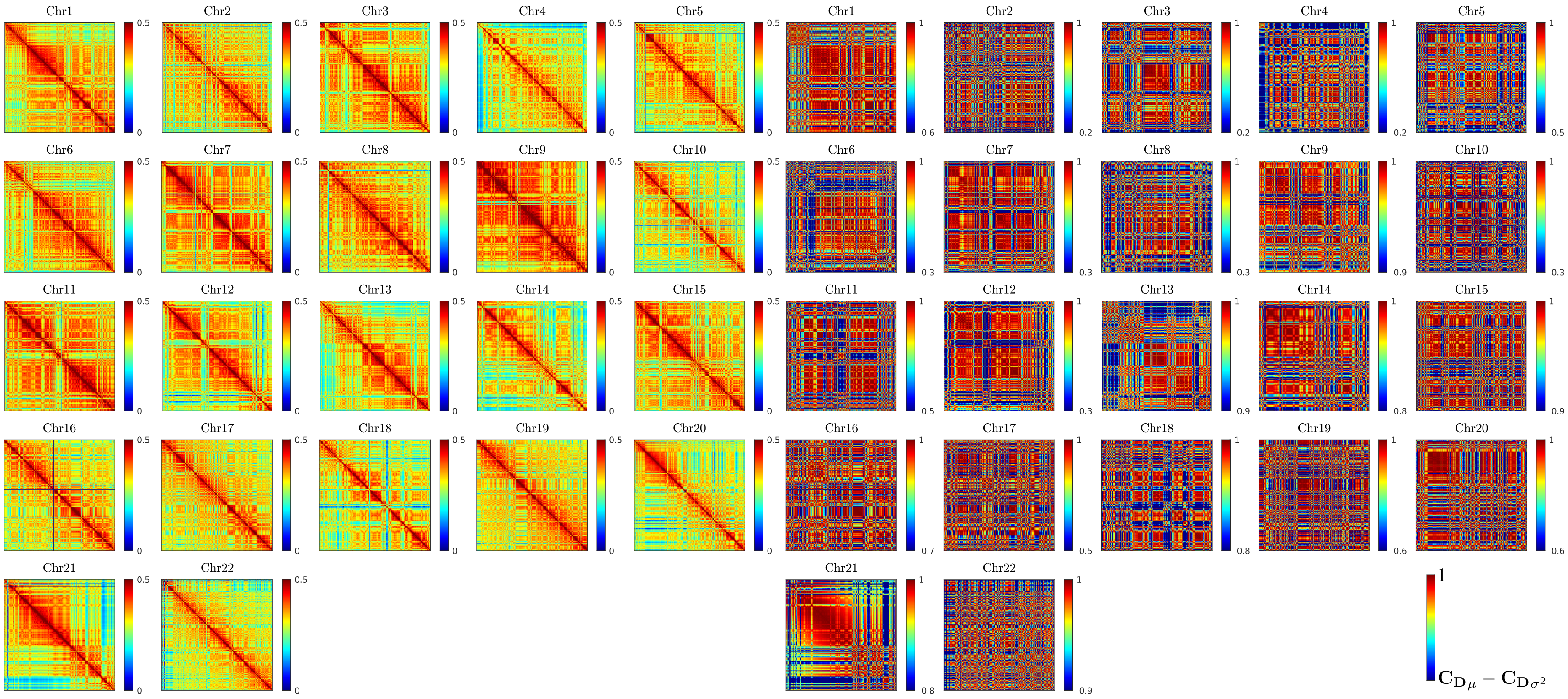


Figure S9: **(left) DNA-DDA contact maps D for GM12878.** Color scale goes from 0 to (right) DNA-DDA pearson contact maps $\mathbf{C_D}$ for GM12878. Color scale goes from mean-variance ($\mathbf{C_{D}}_{\mu} - \mathbf{C_{D}}_{\sigma^2}$) to max

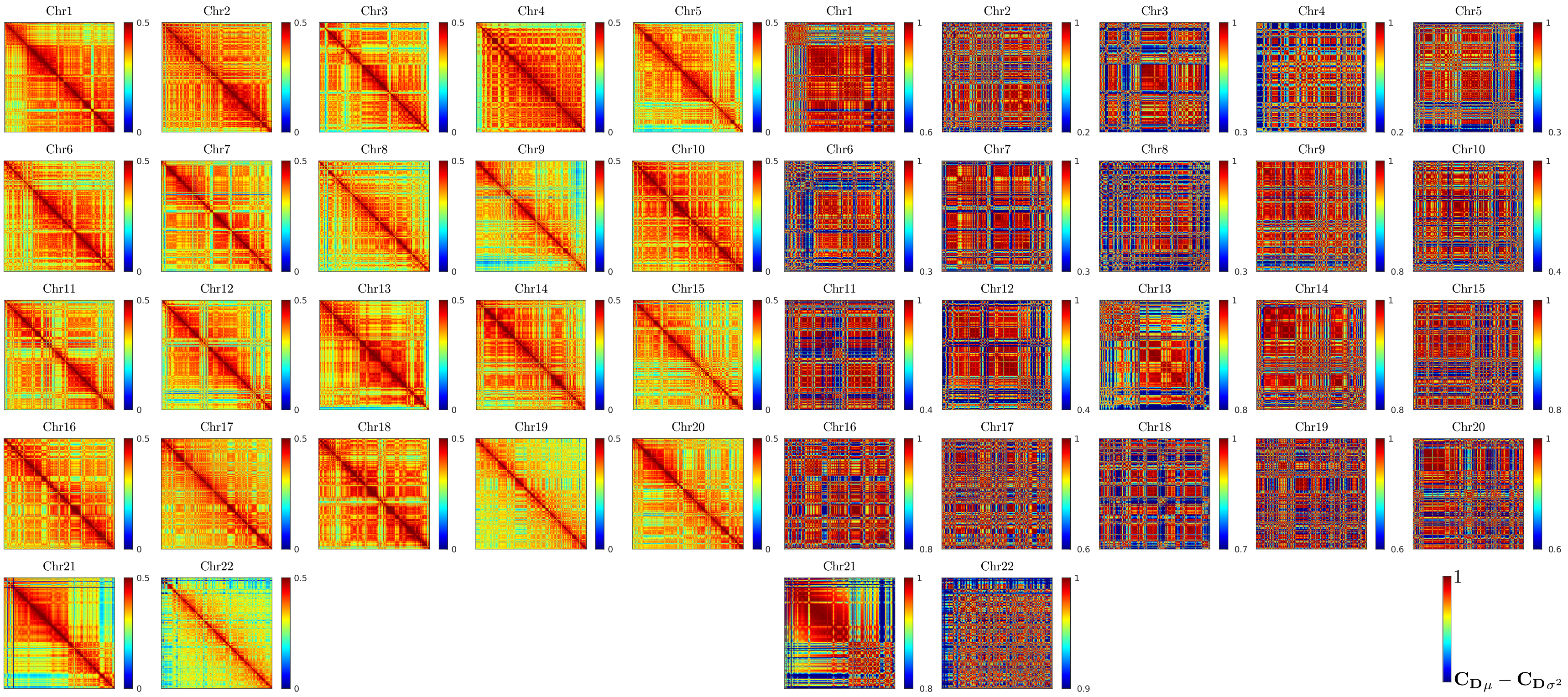


Figure S10: **(left) DNA-DDA contact maps D for K562.** Color scale goes from 0 to (right) DNA-DDA pearson contact maps $\mathbf{C_D}$ for K562. Color scale goes from mean-variance ($\mathbf{C}_{\mathbf{D}_\mu} - \mathbf{C}_{\mathbf{D}_{\sigma^2}}$) to max

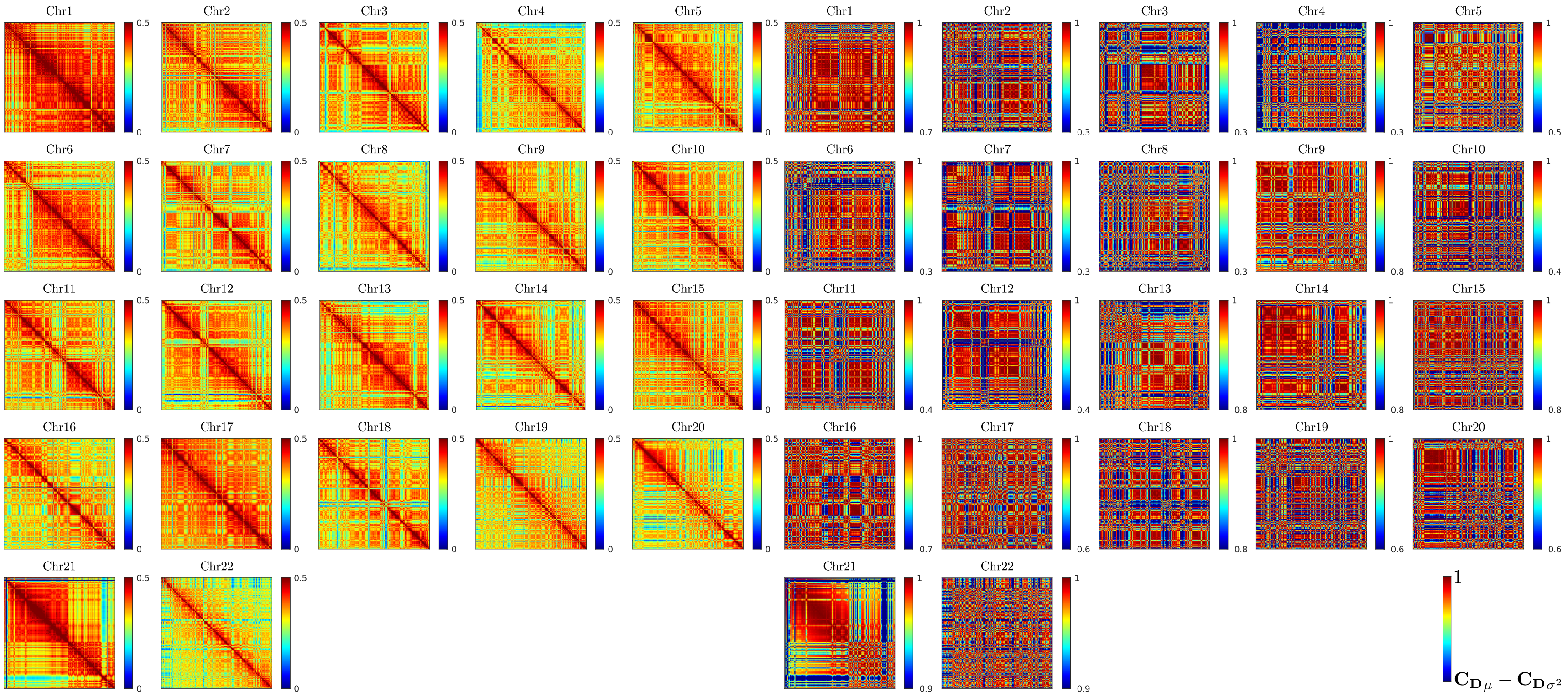


Figure S11: **(left) DNA-DDA contact maps D for IMR90.** Color scale goes from 0 to (right) DNA-DDA pearson contact maps $\mathbf{C_D}$ for IMR90. Color scale goes from mean-variance ($\mathbf{C}_{\mathbf{D}_\mu} - \mathbf{C}_{\mathbf{D}_{\sigma^2}}$) to max

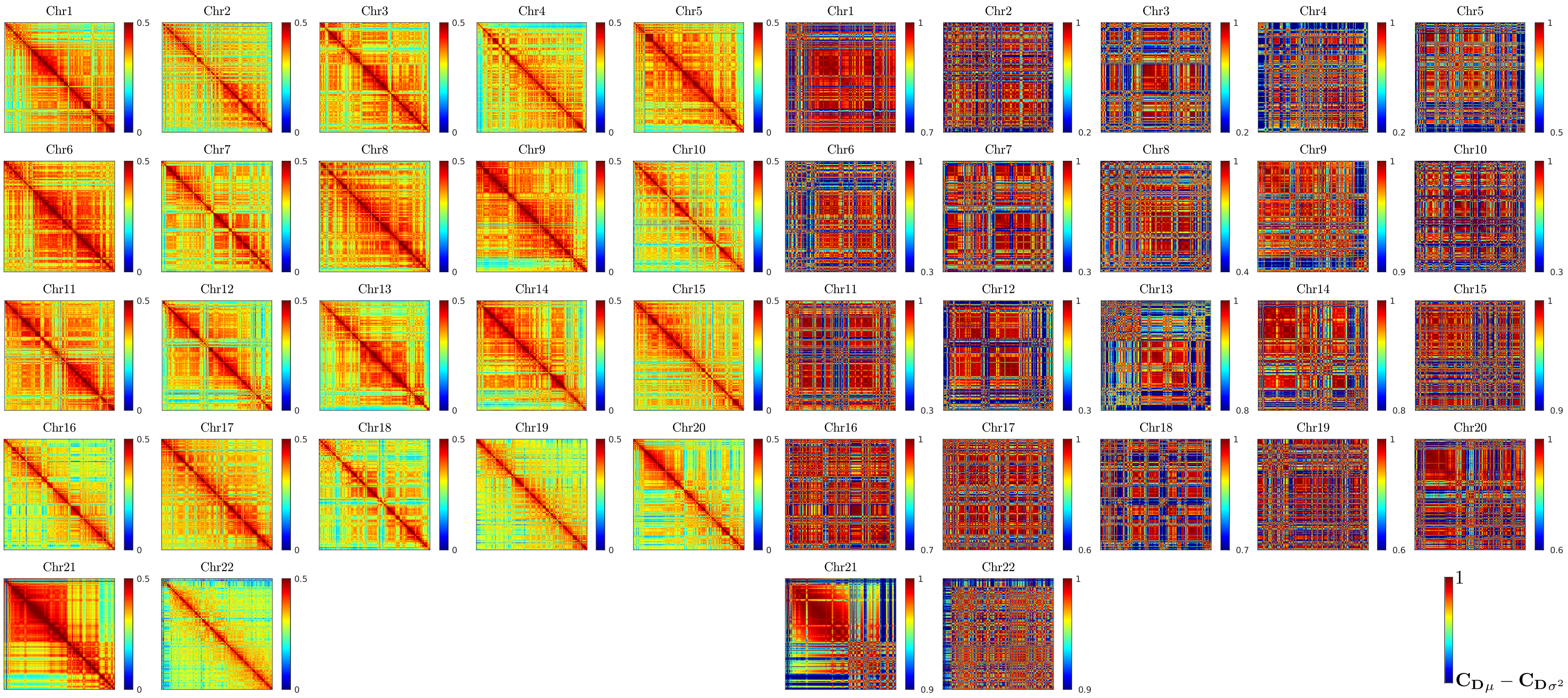


Figure S12: **(left) DNA-DDA contact maps D for hESC.** Color scale goes from 0 to (right) DNA-DDA pearson contact maps $\mathbf{C_D}$ for hESC. Color scale goes from mean-variance ($\mathbf{C}_{\mathbf{D}_\mu} - \mathbf{C}_{\mathbf{D}_{\sigma^2}}$) to max

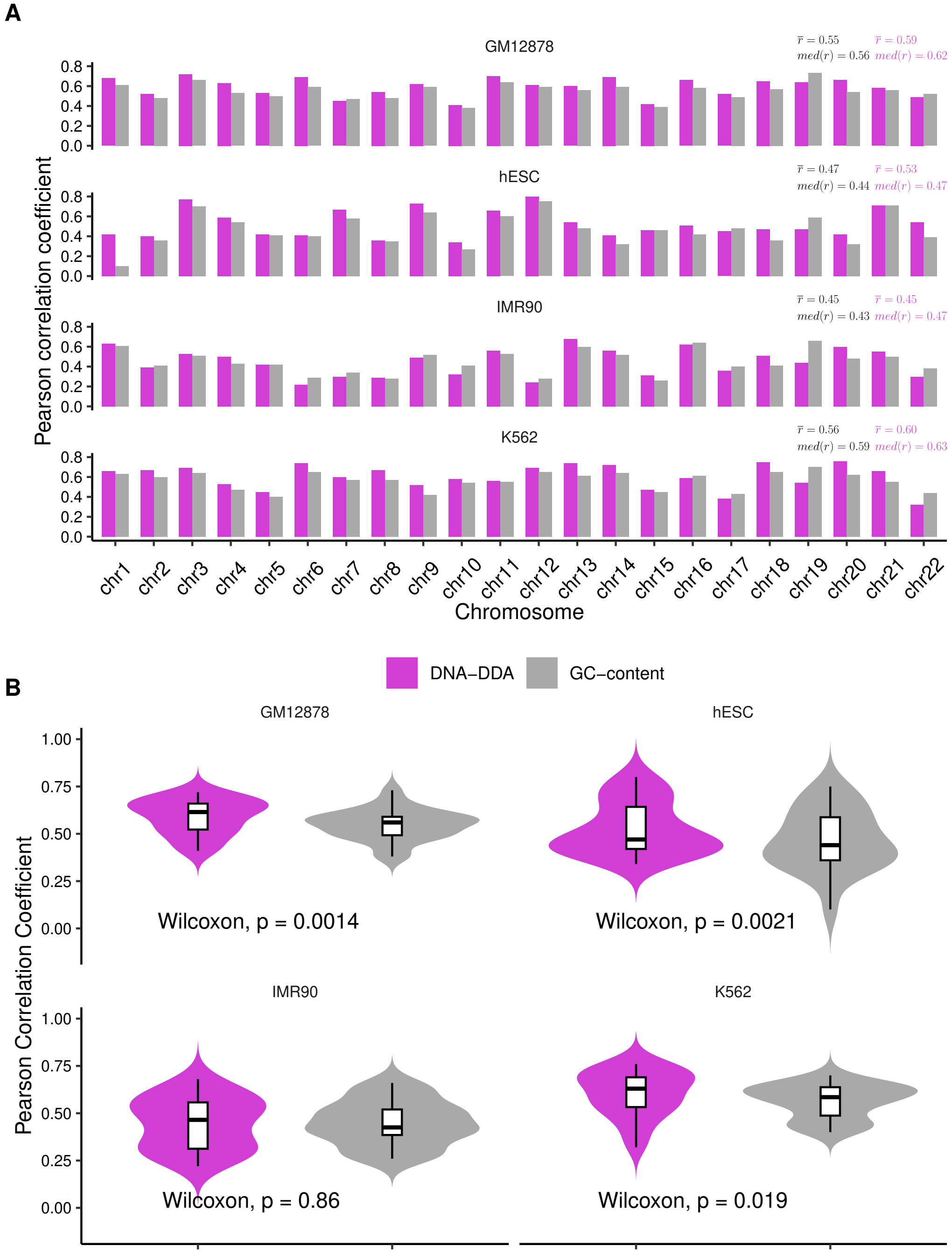


Figure S13: **DNA-DDA predictions are not only driven by the GC-content.** Pearson correlation coefficients between $PC_{\text{DNA-DDA}}$ and $PC_{\text{Hi-C}}$ compared to Pearson correlation coefficients between average GC-content and $PC_{\text{Hi-C}}$. Average GC-content was computed for the same consecutive non-overlapping 100kb-long genomic bins used for predicting compartments with DNA-DDA. A) Per chromosome. B) Per cell line. The P-value corresponds to the Wilcoxon signed-rank test. DNA-DDA outperforms the naïve GC-content-based compartment prediction.

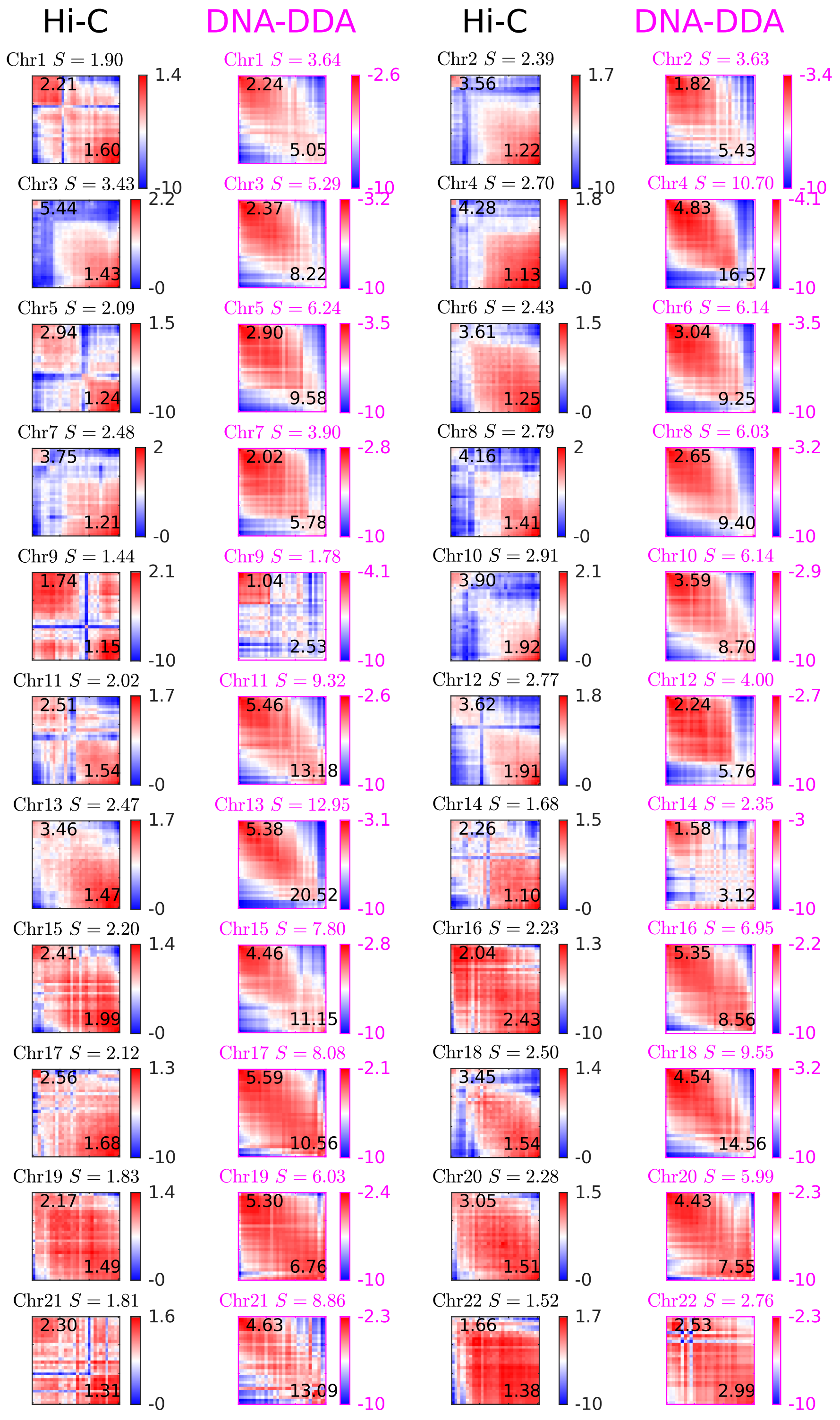


Figure S14: **Saddle plot analysis in K562.** Saddle plots derived from observed/expected Hi-C (black) and DNA-DDA (magenta) matrices. Colors indicate log2 of interaction values given by $HiC_{OE}$ and $\mathbf{D}$ respectively. Compartment strengths $AA$, $BB$, $AB$ and $BA$ are shown in each corner and correspond to the highest ($AA$, $BB$) and lowest ($AB$, $BA$) 25% of the PC values. The overall saddle strength $S$ is then calculated as $\frac{AA+BB}{AB+BA}$.

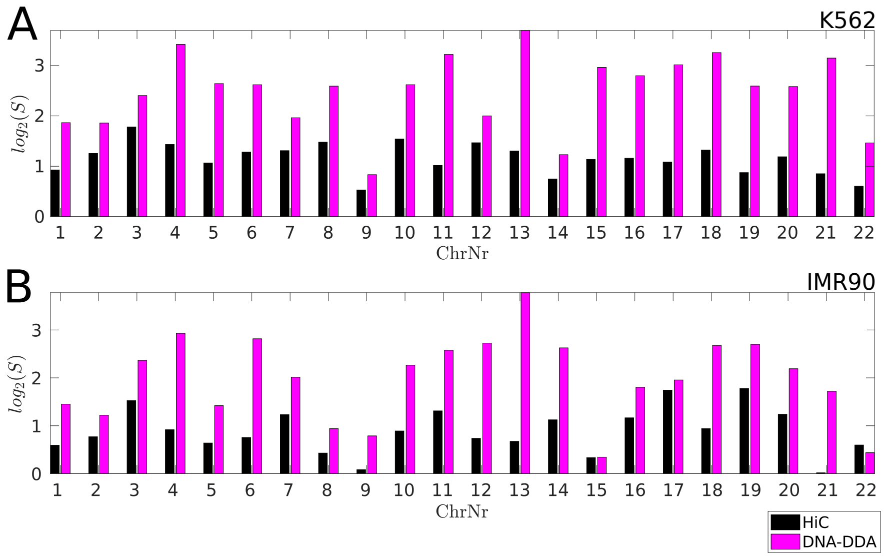


Figure S15: **DNA-DDA predicts overall stronger saddle strengths S.** Barplots of log2 saddle strengths across all chromosomes in A) K562 and B) IMR90.

### Tables

Table S1: `--filterThreshold` parameter for normalizing contact matrices

| Chr | GM12878 | | K562 | | IMR90 | | hESC | |
|---|---|---|---|---|---|---|---|---|
| | $Z_{min}$ | $Z_{max}$ | $Z_{min}$ | $Z_{max}$ | $Z_{min}$ | $Z_{max}$ | $Z_{min}$ | $Z_{max}$ |
| 1 | -2.0 | 2.4 | -1.5 | 3.0 | -0.7 | 2.0 | -0.6 | 2.4 |
| 2 | -2.0 | 2.6 | -1.6 | 3.0 | -0.8 | 2.0 | -0.7 | 2.2 |
| 3 | -2.0 | 2.6 | -1.5 | 4.0 | -0.8 | 2.1 | -0.7 | 2.2 |
| 4 | -2.0 | 3.0 | -1.8 | 3.4 | -0.9 | 2.1 | -0.8 | 2.4 |
| 5 | -2.0 | 2.8 | -1.7 | 3.0 | -0.8 | 2.1 | -0.7 | 2.4 |
| 6 | -1.9 | 2.6 | -1.5 | 3.0 | -0.8 | 2.0 | -0.8 | 2.4 |
| 7 | -1.9 | 2.6 | -1.6 | 3.0 | -0.7 | 1.9 | -0.6 | 2.2 |
| 8 | -1.9 | 2.6 | -1.6 | 3.0 | -0.8 | 2.0 | -0.7 | 2.4 |
| 9 | -1.6 | 2.2 | -1.1 | 3.0 | -0.7 | 1.9 | -0.6 | 2.1 |
| 10 | -2.0 | 2.8 | -1.8 | 4.0 | -0.8 | 2.0 | -0.7 | 2.4 |
| 11 | -1.9 | 2.6 | -1.4 | 2.6 | -0.7 | 2.0 | -0.7 | 2.2 |
| 12 | -2.0 | 2.6 | -1.6 | 3.0 | -0.8 | 2.0 | -0.6 | 2.2 |
| 13 | -1.8 | 2.6 | -1.6 | 3.0 | -0.9 | 2.0 | -0.8 | 2.4 |
| 14 | -1.8 | 2.9 | -1.6 | 3.0 | -0.9 | 2.0 | -0.7 | 2.2 |
| 15 | -2.0 | 2.8 | -1.8 | 2.8 | -0.7 | 1.9 | -0.6 | 2.2 |
| 16 | -2.0 | 2.2 | -1.7 | 2.0 | -0.6 | 2.0 | -0.5 | 2.4 |
| 17 | -2.0 | 2.1 | -1.6 | 2.0 | -0.5 | 2.0 | -0.4 | 2.6 |
| 18 | -1.9 | 2.6 | -1.4 | 3.2 | -0.8 | 2.0 | -0.7 | 2.2 |
| 19 | -2.4 | 2.0 | -2.1 | 2.0 | -0.5 | 2.0 | -0.4 | 2.8 |
| 20 | -2.2 | 2.0 | -1.4 | 2.4 | -0.7 | 1.9 | -0.5 | 2.1 |
| 21 | -1.4 | 2.4 | -1.2 | 3.0 | -0.7 | 1.9 | -0.5 | 2.1 |
| 22 | -1.8 | 2.0 | -1.9 | 2.0 | -0.5 | 2.0 | -0.4 | 2.0 |

Table S2: Parameters used to call the `C` executable implementation of DDA. This applies an SVD algorithm to obtain model features and the least squares error ($\{a_1, a_2, a_3, \rho\}$) for an input sequence $x$ and model 2 in Supplementary Table S4

| variable | value |
|---|---|
| `TAU_FILE` | `DELAY_FILE` |
| `DATA_FN` | `RandomWalk_Chr1.ascii` |
| `OUT_FN` | `DDA_OUT/Chr1_start_end_bin#.ascii` |
| `WL` | 99942 |
| `WS` | 99942 |
| `SELECT` | `ST CT 0 0` |
| `CT_CH_list` | $b_s\ b_{s+1}\ b_s\ b_{s+2} \ldots b_s\ b_{N_{bins}}$ |

- `DELAY_FILE` ... file with 2 columns containing values of delay values $\tau_1, \tau_2$
- `RandomWalk_Chr1.ascii` ... file containing $N_{bins}$ input sequences of length 100kbp ($N_{bins} = 2490$ for chromosome 1)
- `OUT_FN` ... file to store outputs of DDA
- `WL` ... window length from which $\{a_1, a_2, a_3, \rho\}$ are extracted from input data
- `WS` ... window shift
- `SELECT` ... for ST DDA: `1 0 0 0`; for CT DDA: `0 1 0 0`;
- $b_s\ b_{s+1}\ b_s\ b_{s+2} \ldots b_s\ b_{N_{bins}}$ ... list of bins to CT

Table S3: Structure selection results for DNA-DDA. Performance is given for the delay pair $\tau_1, \tau_2$ that resulted in the highest correlation coefficient $r_{PC}$ between $PC_{\text{DNA-DDA}}$ and $PC_{\text{Hi-C}}$ on chr22:16200000:36200001 (Figure S2).

| Cell type | $\tau_1$ | $\tau_2$ | $r_{PC_{Chr22SS}}$ | $r_{PC_{Chr22ALL}}$ |
|---|---|---|---|---|
| GM12878 | 3 | 6 | 0.49 | 0.49 |
| K562 | 11 | 50 | 0.42 | 0.32 |
| hESC | 2 | 4 | 0.21 | 0.53 |
| IMR90 | 1 | 36 | 0.35 | 0.30 |

Table S4: All possible three term DDA models up to cubic nonlinearity and two delays. A checkmark "✓" denotes term $a_i$ in Eq. to be nonzero. Symmetric models are labeled with "S" and single delay models with "1". All others have two non-interchangable delays.

| ModelNr | $x_1$ | $x_2$ | $x_1^2$ | $x_1x_2$ | $x_2^2$ | $x_1^3$ | $x_1^2x_2$ | $x_1x_2^2$ | $x_2^3$ | model type |
|---|---|---|---|---|---|---|---|---|---|---|
| 1 | ✓ | ✓ | ✓ | 0 | 0 | 0 | 0 | 0 | 0 | |
| 2 | ✓ | ✓ | 0 | ✓ | 0 | 0 | 0 | 0 | 0 | S |
| 3 | ✓ | ✓ | 0 | 0 | 0 | ✓ | 0 | 0 | 0 | |
| 4 | ✓ | ✓ | 0 | 0 | 0 | 0 | ✓ | 0 | 0 | |
| 5 | ✓ | 0 | ✓ | ✓ | 0 | 0 | 0 | 0 | 0 | |
| 6 | ✓ | 0 | ✓ | 0 | ✓ | 0 | 0 | 0 | 0 | |
| 7 | ✓ | 0 | ✓ | 0 | 0 | ✓ | 0 | 0 | 0 | 1 |
| 8 | ✓ | 0 | ✓ | 0 | 0 | 0 | ✓ | 0 | 0 | |
| 9 | ✓ | 0 | ✓ | 0 | 0 | 0 | 0 | ✓ | 0 | |
| 10 | ✓ | 0 | ✓ | 0 | 0 | 0 | 0 | 0 | ✓ | |
| 11 | ✓ | 0 | 0 | ✓ | ✓ | 0 | 0 | 0 | 0 | |
| 12 | ✓ | 0 | 0 | ✓ | 0 | ✓ | 0 | 0 | 0 | |
| 13 | ✓ | 0 | 0 | ✓ | 0 | 0 | ✓ | 0 | 0 | |
| 14 | ✓ | 0 | 0 | ✓ | 0 | 0 | 0 | ✓ | 0 | |
| 15 | ✓ | 0 | 0 | ✓ | 0 | 0 | 0 | 0 | ✓ | |
| 16 | ✓ | 0 | 0 | 0 | ✓ | ✓ | 0 | 0 | 0 | |
| 17 | ✓ | 0 | 0 | 0 | ✓ | 0 | ✓ | 0 | 0 | |
| 18 | ✓ | 0 | 0 | 0 | ✓ | 0 | 0 | ✓ | 0 | |
| 19 | ✓ | 0 | 0 | 0 | ✓ | 0 | 0 | 0 | ✓ | |
| 20 | ✓ | 0 | 0 | 0 | 0 | ✓ | ✓ | 0 | 0 | |
| 21 | ✓ | 0 | 0 | 0 | 0 | ✓ | 0 | ✓ | 0 | |
| 22 | ✓ | 0 | 0 | 0 | 0 | ✓ | 0 | 0 | ✓ | |
| 23 | ✓ | 0 | 0 | 0 | 0 | 0 | ✓ | ✓ | 0 | |
| 24 | ✓ | 0 | 0 | 0 | 0 | 0 | ✓ | 0 | ✓ | |
| 25 | ✓ | 0 | 0 | 0 | 0 | 0 | 0 | ✓ | ✓ | |
| 26 | 0 | 0 | ✓ | ✓ | ✓ | 0 | 0 | 0 | 0 | S |
| 27 | 0 | 0 | ✓ | ✓ | 0 | ✓ | 0 | 0 | 0 | |
| 28 | 0 | 0 | ✓ | ✓ | 0 | 0 | ✓ | 0 | 0 | |
| 29 | 0 | 0 | ✓ | ✓ | 0 | 0 | 0 | ✓ | 0 | |
| 30 | 0 | 0 | ✓ | ✓ | 0 | 0 | 0 | 0 | ✓ | |
| 31 | 0 | 0 | ✓ | 0 | ✓ | ✓ | 0 | 0 | 0 | |
| 32 | 0 | 0 | ✓ | 0 | ✓ | 0 | ✓ | 0 | 0 | |
| 33 | 0 | 0 | ✓ | 0 | 0 | ✓ | ✓ | 0 | 0 | |
| 34 | 0 | 0 | ✓ | 0 | 0 | ✓ | 0 | ✓ | 0 | |
| 35 | 0 | 0 | ✓ | 0 | 0 | ✓ | 0 | 0 | ✓ | |
| 36 | 0 | 0 | ✓ | 0 | 0 | 0 | ✓ | ✓ | 0 | |
| 37 | 0 | 0 | ✓ | 0 | 0 | 0 | ✓ | 0 | ✓ | |
| 38 | 0 | 0 | ✓ | 0 | 0 | 0 | 0 | ✓ | ✓ | |
| 39 | 0 | 0 | 0 | ✓ | 0 | ✓ | ✓ | 0 | 0 | |
| 30 | 0 | 0 | 0 | ✓ | 0 | ✓ | 0 | ✓ | 0 | |
| 41 | 0 | 0 | 0 | ✓ | 0 | ✓ | 0 | 0 | ✓ | S |
| 42 | 0 | 0 | 0 | ✓ | 0 | 0 | ✓ | ✓ | 0 | S |
| 43 | 0 | 0 | 0 | 0 | 0 | ✓ | ✓ | ✓ | 0 | |
| 44 | 0 | 0 | 0 | 0 | 0 | ✓ | ✓ | 0 | ✓ | |

Table S5: Omitted genomic bins. Includes centromeres from UCSD genome browser and low coverage bins of HiC contact maps in GM12878 (less than 10% of overall interaction probability).

| | |
|---|---|
| chr1 | 1:5, 1219:1250, 1253:1431, 1442:1444, 1447:1448, 1463, 1488 |
| chr2 | 895:895, 906:914, 921:941, 944, 1099, 1105 |
| chr3 | 907:937 |
| chr4 | 497:518 |
| chr5 | 464:501, 698:699 |
| chr6 | 585:602 |
| chr7 | 581:617, 621:621 |
| chr8 | 1, 440:459 |
| chr9 | 393, 400, 433:605, 607, 609, 611, 623, 631:632, 635:635, 643, 647, 651:652, 659, 665:666, 680:682 |
| chr10 | 396:417, 479 |
| chr11 | 1, 510:545 |
| chr12 | 347:372 |
| chr13 | 1:181 |
| chr14 | 1:190, 196 |
| chr15 | 1:220 |
| chr16 | 185, 332:333, 344, 363:463 |
| chr17 | 1, 228:267 |
| chr18 | 154:209 |
| chr19 | 244:272 |
| chr20 | 264:301 |
| chr21 | 1:50, 55:60, 63:63, 67:68, 71:72, 75:77, 82:82, 84:84, 95:95, 108:130 |
| chr22 | 1:105, 109, 112, 129:152 |

Table S6: Omitted genomic bins. Includes centromeres from UCSD genome browser and low coverage bins of HiC contact maps in K562 (less than 10% of overall interaction probability).

| | |
|---|---|
| chr1 | 1:5, 1219:1250, 1253:1431, 1442:1444, 1447:1449, 1463, 1488 |
| chr2 | 895, 906:914, 921:941, 944:945, 1099, 1105:1106 |
| chr3 | 907:937 |
| chr4 | 497:518 |
| chr5 | 464:501, 698:699, 703, 707 |
| chr6 | 1, 585:602, |
| chr7 | 581:617, 621 |
| chr8 | 1, 440:459, |
| chr9 | 209:265, 287:316, 393, 400, 405, 407, 427, 433:605, 607:611, 615, 622:627, 631:632, 635, 643, 647:648, 651:652, 659, 665:666, 673, 676, 680:682 |
| chr10 | 396:417, 1020 |
| chr11 | 1, 510:545 |
| chr12 | 347:372 |
| chr13 | 1:181 |
| chr14 | 1:183, 189, 196 |
| chr15 | 1:198 |
| chr16 | 185:185, 328, 332:333, 344:345, 363:463 |
| chr17 | 1, 228:267 |
| chr18 | 154:209 |
| chr19 | 1, 244:272 |
| chr20 | 264:301 |
| chr21 | 1:50, 55:60, 63, 67:68, 71:72, 75:77, 81:84, 95:95, 108:130 |
| chr22 | 1:106, 109, 129:152 |

Table S7: Omitted genomic bins. Includes centromeres from UCSD genome browser and low coverage bins of HiC contact maps in IMR90 (less than 10% of overall interaction probability).

| | |
|---|---|
| chr1 | 1:7, 1203, 1207:1209, 1214, 1217:1250, 1253:1435, 1440:1444, 1447:1452, 1454, 1462:1464, 1486:1489, 1493:1496 |
| chr2 | 870, 878, 895:896, 898, 905:914, 921:941, 944:945, 975, 1099:1101, 1104:1106, 1306 |
| chr3 | 907:937 |
| chr4 | 93, 495, 497:518 |
| chr5 | 464:501, 697:710, 712:713 |
| chr6 | 1, 585:602 |
| chr7 | 581:617, 731:732, 749, 752:754, 1437 |
| chr8 | 1:2, 73, 76:78, 80:81, 123, 440:459, 858 |
| chr9 | 393:394, 397:398, 400:401, 403:407, 427, 433:605, 607:615, 619:628, 630:633, 635, 643:644, 646:648, 651:653, 658:668, 672:683 |
| chr10 | 396:417, 420:421, 479, 481 |
| chr11 | 1:2, 510:545, 900 |
| chr12 | 347:372 |
| chr13 | 1:181 |
| chr14 | 1:190, 196 |
| chr15 | 1:222, 224:225, 286:287, 305:306, 325, 843, 845 |
| chr16 | 164, 184:185, 215, 288, 296, 328, 332:335, 344:345, 363:463 |
| chr17 | 1, 192, 228:267, 364:364, 380:382, 464, 466 |
| chr18 | 154:209 |
| chr19 | 1:2, 244:272 |
| chr20 | 264:301, 312 |
| chr21 | 1:52, 55:63, 65:72, 75:79, 81:85, 87, 95, 103, 108:130, 138, 435, 442 |
| chr22 | 1:106, 109, 112, 118, 123, 129:153, 164, 183:188, 213:214 |

Table S8: Omitted genomic bins. Includes centromeres from UCSD genome browser and low coverage bins of HiC contact maps in hESC (less than 10% of overall interaction probability).

| | |
|---|---|
| chr1 | 1:8, 1203, 1207:1209, 1213:1214, 1217:1250, 1253:1431, 1434:1435, 1438:1452, 1454:1455, 1462:1464, 1486:1489, 1492:1496 |
| chr2 | 870, 878, 894:896, 898, 905:914, 921:941, 944:945, 975, 1099:1101, 1104:1106, 1306 |
| chr3 | 907:937, 1629 |
| chr4 | 495, 497:518 |
| chr5 | 464:501, 697:710, 712:713 |
| chr6 | 1, 585:602 |
| chr7 | 581:617, 621, 731:733, 749:750, 752:754, 1026, 1437 |
| chr8 | 1:2, 73, 76:78, 81, 440:459, 858 |
| chr9 | 393, 397:407, 412, 415:417, 419, 427, 433:605, 607:616, 618:628, 630:636, 643:644, 646:649, 651:656, 658:667, 672:683 |
| chr10 | 396:417, 479 |
| chr11 | 1:2, 510:545 |
| chr12 | 347:372 |
| chr13 | 1:181, 1118 |
| chr14 | 1:186, 188:190, 194:196 |
| chr15 | 1:198, 201, 206, 208:209, 211:212, 214:222, 224:225, 233, 305:306, 325, 843, 845 |
| chr16 | 148, 164, 184, 215, 288, 296, 303, 328, 331:335, 344:345, 363:463 |
| chr17 | 1, 192:192, 228:267, 364, 380:382, 464 |
| chr18 | 154:209 |
| chr19 | 1:2, 244:272 |
| chr20 | 264:301, 312 |
| chr21 | 1:52, 55:63, 65:72, 75:79, 81:84, 86:87, 95, 103, 108:130, 138, 435:435, 442:442 |
| chr22 | 1:107, 109, 112, 129:156, 158:159, 183:188, 214 |

Table S9: Principal components used to determine A/B compartments from Hi-C and DNA-DDA matrix respectively

| Chr | GM12878 | | K562 | | IMR90 | | hESC | |
|---|---|---|---|---|---|---|---|---|
| | Hi-C | DNA-DDA | Hi-C | DDA | Hi-C | DDA | Hi-C | DDA |
| 1 | 1 | 1 | 1 | 1 | 2 | 1 | 1 | 4 |
| 2 | 1 | 1 | 1 | 1 | 2 | 1 | 1 | 1 |
| 3 | 1 | 1 | 2 | 1 | 2 | 1 | 3 | 1 |
| 4 | 1 | 1 | 2 | 1 | 2 | 1 | 4 | 1 |
| 5 | 1 | 1 | 2 | 1 | 2 | 1 | 3 | 1 |
| 6 | 1 | 1 | 1 | 1 | 2 | 2 | 4 | 1 |
| 7 | 1 | 1 | 1 | 1 | 2 | 1 | 2 | 1 |
| 8 | 1 | 1 | 1 | 1 | 2 | 1 | 4 | 1 |
| 9 | 1 | 2 | 2 | 1 | 2 | 2 | 2 | 1 |
| 10 | 1 | 1 | 1 | 1 | 2 | 2 | 3 | 1 |
| 11 | 1 | 2 | 1 | 1 | 2 | 2 | 2 | 1 |
| 12 | 1 | 1 | 1 | 1 | 1 | 2 | 2 | 1 |
| 13 | 1 | 1 | 1 | 1 | 1 | 1 | 4 | 1 |
| 14 | 1 | 2 | 1 | 1 | 1 | 2 | 3 | 1 |
| 15 | 1 | 1 | 1 | 1 | 1 | 1 | 2 | 1 |
| 16 | 1 | 2 | 1 | 1 | 1 | 2 | 4 | 1 |
| 17 | 1 | 2 | 1 | 1 | 1 | 2 | 2 | 1 |
| 18 | 1 | 1 | 1 | 1 | 1 | 2 | 4 | 1 |
| 19 | 1 | 2 | 1 | 1 | 1 | 1 | 1 | 2 |
| 20 | 1 | 2 | 1 | 1 | 1 | 2 | 2 | 1 |
| 21 | 1 | 2 | 1 | 2 | 1 | 2 | 2 | 2 |
| 22 | 1 | 2 | 1 | 2 | 1 | 2 | 2 | 4 |

Table S10: Overall datasets used by comparison methods

| | | |
|---|---|---|
| SACSANN | GSE35156 | mESC,hESC |
| | GSE96107 | mESC,NPC,CN |
| | GSE59027 | mESC,NPC,CN |
| ABCNet | GSE35156 | hESC |
| | GSE48592,GSE63525 | EBV |
| | GSE43070,GSE63525 | IMR90 |
| | GSE31388 | Dnase EBV |
| | GSE31263 | Dnase IMR90 |
| | GSE53261 | Hi-C meth fibroblast |
| | http://www.roadmapepigenomics.org/ | Hi-C meth EBV |
| Orca | 4DNFI9GMP2J8 | H1ESC |
| | 4DNFI643OYP9 | HFF |
| | 4DNFILP99QJS | cohesin-depleted HCT116 |
| DNA-DDA | GSE63525 | GM12878,K562 |
| | GSE35156 | hESC,IMR90 |

Table S11: Regions used for Orca comparison.

| | |
|---|---|
| 1 | chr9:5504001-37504000 |
| 2 | chr9:69888001-101888000 |
| 3 | chr9:104960001-136960000 |
